# Potassium dependent structural changes in the selectivity filter of HERG potassium channels

**DOI:** 10.1101/2023.12.14.571769

**Authors:** Carus H.Y. Lau, Emelie Flood, Mark J. Hunter, Billy J Williams-Noonan, Karen M. Corbett, Chai-Ann Ng, James C. Bouwer, Alastair G. Stewart, Eduardo Perozo, Toby W. Allen, Jamie I. Vandenberg

**Affiliations:** Mark Cowley Lidwill Research Program, Victor Chang Cardiac Research Institute; Darlinghurst, NSW, Australia; School of Clinical Medicine, UNSW, Sydney, NSW, Australia; School of Science, RMIT University; Melbourne, Vic, Australia; Molecular Horizons and School of Chemistry and Molecular Bioscience, University of Wollongong, NSW, Australia; Computational and Structural Biology Division, Victor Chang Cardiac Research Institute; Darlinghurst, NSW, Australia; Department of Biochemistry and Molecular Biology, The University of Chicago, IL, USA

## Abstract

The fine tuning of biological electrical signaling is mediated by variations in the rates of opening and closing of gates that control ion flux through different ion channels. Human ether-a-go-go related gene (HERG) potassium channels have uniquely rapid inactivation kinetics which are critical to the role they play in regulating cardiac electrical activity. Here, we have exploited the K^+^ sensitivity of HERG inactivation to determine structures of both a conductive and non-conductive selectivity filter structure of HERG. We propose that inactivation is the result of a high propensity for flipping of the selectivity filter valine carbonyl oxygens. Molecular dynamics simulations point to a low energy barrier, and hence rapid kinetics, for flipping of the valine 625 carbonyl oxygens facilitated by a previously unrecognized interaction between S620 and Y616 that stabilizes the transition state between conducting and non-conducting structures. Our model represents a new mechanism by which ion channels fine tune their activity that explains the uniquely rapid inactivation kinetics of HERG.

**Highlights:** Structures of a conductive and non-conductive HERG selectivity filter have been determined.

Reduced potassium causes flipping of selectivity filter valine carbonyl oxygens.

The sidechain of S620 on the pore helix coordinates distinct sets of interactions between conductive, non-conductive, and transition states.

## Introduction

Ion channels may exist in one of three main conformational states: closed, open, or inactivated. Inactivation describes the process whereby ion channels enter a non-conducting state in the presence of a prolonged activating stimulus. Differences in rates of inactivation of ion channels are a major determinant of their specific physiological functions^1^. In HERG K+ channels, the combination of the rapidity, voltage dependence, and K^+^ sensitivity of inactivation is critical for regulating cardiac repolarization and the response to ectopic beats^2–4^. Furthermore, both hypokalaemia^5^ and mutations that affect inactivation gating^6,7^ increase the risk of cardiac arrhythmias. Despite the clinical importance of HERG potassium channels, we still do not understand the structural basis for how they inactivate.

One of the major mechanisms underlying inactivation in K^+^ channels is conformational changes in the selectivity filter resulting in loss of conduction^1^. There is very high sequence conservation in the selectivity filter of different classes of potassium channels ^1^ with subtle sequence differences in the selectivity filter and immediately behind the filter contributing to the differences in the rates of inactivation of different K^+^ channels^8,9^. For example, the rate of inactivation of HERG K^+^ channels is approximately two orders of magnitude faster than inactivation of the bacterial KcsA K^+^ channel and approximately three orders of magnitude faster than c-type inactivation of the eukaryotic Shaker K^+^ channel^4^. Why inactivation in HERG is so fast remains unresolved.

One approach that has been used to probe the structural basis of K^+^ channel selectivity filter gating is to determine structures in the presence of different K^+^ concentrations^10–12^. In the bacterial KcsA K^+^ channel, the most extensively studied K^+^ channel structure, the selectivity filter has a cylindrical profile in the presence of 300 mM K^+^ with five K^+^ ion binding sites, denoted S0 - S4^10,13^. In the presence of 3 mM K^+^, the KcsA selectivity filter has a constriction at the level of G77 Cα^10,11^ and rotation of the valine (V76) carbonyl oxygens to point ∼90° away from the central axis^10^. These changes are also observed in mutant channels with a forced open activation gate^14^ and recapitulated in MD simulations^11,15–17^. Thus, a consensus has emerged that the inactivated state of KcsA has an hourglass shaped selectivity filter with the constriction at the level of G77. In contrast to KcsA, a recent study of heterodimeric K_2P.1_ channels showed that <1 mM K^+^ causes an asymmetrical constriction and dilatation of the filter ^12^. Furthermore, in the Shaker family of voltage gated K^+^ channels, mutations that enhance inactivation result in dilatation of the extracellular half of the filter with no constriction at the central glycine^18–21^. Thus, whilst the structures of the selectivity filter in the conductive state of different classes of potassium channels are similar, it is likely that they have adopted different mechanisms for conversion to a non-conducting state.

Cryo-EM studies of WT HERG and the inactivation-deficient S631A HERG channels, in the presence of 300 mM K^+^, revealed an ∼10° rotation of F627, the selectivity filter phenylalanine in the selectivity filter^22^. The authors suggested that such a subtle structural change was consistent with the rapidity of inactivation in HERG^22^. Conversely, Molecular dynamics (MD) simulation studies of the WT HERG structure suggested that the transition from conducting to non-conducting selectivity filter structures could involve asymmetric buckling of the selectivity filter^23,24^, and/or dilation of the upper filter^23,25^. An alternative explanation for the similarity of the WT and S631A HERG structures is that they both represent subtly different conductive state structures^23,26,27^. Thus, the structural basis for HERG inactivation remains unresolved. Here, to explore the structural basis for the rapidity of HERG inactivation, we have used cryo-EM to investigate the potassium dependent structural changes in the selectivity filter and molecular dynamics (MD) simulations to explore how K^+^ dependent rearrangements of the selectivity filter can occur so rapidly.

## Results

### Structures of HERG in high-K^+^ and low-K^+^ conditions

We used a WT HERG construct with the predicted unstructured regions spanning residues 140-380 and 870-1006 deleted^22^. Structures of WT HERG incubated with 300 mM K^+^ (high-K) or 3 mM K^+^ (low-K) were initially solved with C1 symmetry. However, as there was no substantial asymmetry between subunits (Supplementary Fig **S1**), we solved the structures using C4 symmetry to improve the overall resolution (Fig 1**A**). The global resolution was 3.3 Å for high-K^+^ and 3.0 Å for low-K^+^ (Fig **1B**, Supplementary Fig **S1**, Supplementary Table **S1**), with the resolution in the selectivity filter region reaching 3.1 Å (high-K) and 2.9 Å (low-K) respectively (Fig **1B**).

**Fig 1:**
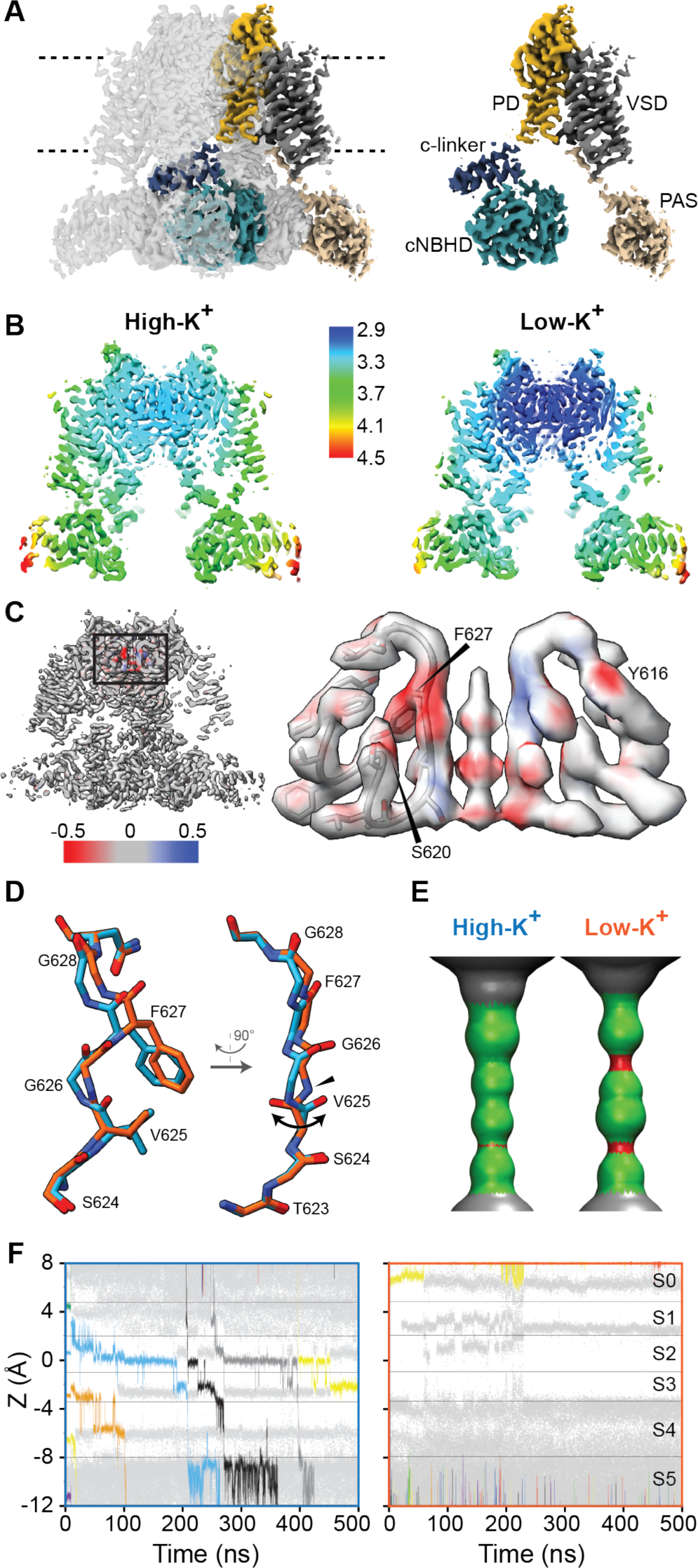
Cryo-EM structure of WT-HERG solved in presence of high-K and low-K. **A**. Cryo-EM map of low-K WT-HERG (contour 6α) with a single subunit of the channel highlighted to illustrate the major domains: pore-domain (PD), voltage sensor domain (VSD), cyclic nucleotide binding homology domain (cNBHD, per-arnt-sim (PAS) domain. **B.** Local resolution of cryo-EM maps solved using C4 symmetry. The resolution range of the transmembrane regions are reported in Supplementary Table 1. **C.** Fractional difference map for high-K and low-K maps. Maps were normalized and low pass filtered to 3.3 Å before fractional difference was calculated using TEMpy:Difference Map tool in CCPEM^29^. Red indicates density only present in the low-K map (density missing in high-K) and blue represents density only present in the high-K map; thresholding 0.0252 (8α). Right panel shows a close-up view of the selectivity filter (high-K, blue and low-K, red). In addition to the selectivity filter residues, there are differences seen for the S620 and Y616 sidechains. **D.** overlay of low-K (orange) and high-K selectivity filter structures, aligned to the pore helix. In the left panel the lateral and upward displacement of the F627/G628 back bone in the low-K structure is apparent as is a small lateral displacement of the S624 sidechain. On the rotated view, the central displacement of the G626 carbonyl, flipping of the V625 carbonyl (bidirectional arrows) and resulting central facing G626 amide (arrowhead) are apparent in the low-K structure. **E.** The high-K structure has a canonical SF HOLE profile, whereas the low-K structure shows dilation in the vicinity of V625 and constrictions at the level of S624 and G626. This is distinct to the single constriction observed in the KcsA low-K filter at G77 (equivalent to G626, see Supplementary Fig **S5**). **F.** Example MD simulations showing ion conduction events when the V625 carbonyl oxygens point towards the central cavity (to mimic high-K structure) but no conduction is seen when the V625 carbonyl oxygens point outwards (to mimic low-K structure). Ions are shown as coloured lines, water molecules as grey points.

The density maps for low-K and high-K structures are very similar, with the major differences occurring in the selectivity filter region (Fig **1C**). The most obvious density difference corresponds to the sidechain and backbone of F627 with other notable differences in the vicinity of the backbone of G626, sidechain and backbone of V625, sidechain of S620, and sidechain of Y616 (Fig **1C**). There are also differences in the density along the ion conduction pathway, which are discussed in more detail below.

The root mean square deviation (RMSD) for the backbone atoms of the best fit structures for the low-K and high-K maps, aligned on the pore helix, was 1.2 Å in the selectivity filter compared to 0.3 Å in the pore domain helices: S5, S5P, and S6; consistent with the minimal difference in density seen away from the selectivity filter (Supplementary Fig **S2**). In the high-K structure, the selectivity filter carbonyl oxygens for S624, V625, G626, and F627 point towards the central axis (Fig **1D**), as seen in previously reported structures for WT HERG ^22,28^, and shows a cylindrical profile (Fig **1E**) similar to that seen in KcsA, Trek-1, and Shaker, when determined in the presence of high-K^10,12,18^.

The differences seen in the low-K structure relative to the high-K structure include (i) a lateral displacement of the F627 backbone atoms and sidechain (ii) displacement of the G626 carbonyl oxygens towards the central axis, (iii) rotation of the V625 carbonyl oxygens such that they now point away from the central axis (bi-directional arrowhead, Fig **1D**) causing the G626 backbone amide to point towards the central axis (blue arrowhead in Fig **1D**), and (iv) a small lateral displacement of the S624 sidechain. At the resolution of our structures, we cannot see the precise location of backbone carbonyl oxygens. Nevertheless, we can infer that in the low-K structure the V625 carbonyl oxygens are flipped to face away from the central cavity, based on (i) higher probability in the Ramachandran plot (Supplementary Fig **S3**) and (ii) molecular dynamics flexible fitting (MDFF) simulations to the cryo-EM density maps (Supplementary Fig **S4**). As a result of the V625 carbonyl oxygens flipping there is a dilation of the ion conduction pathway in the vicinity of V625 in the low-K HERG structure (Fig **1E**) which is quite distinct to the hourglass profile seen in KcsA low-K (Supplementary Fig **S5).**

In previous MD simulations, it has been shown that the high-K structure can conduct K^+^ ions^23^. To specifically investigate the effect of V625 carbonyl oxygens pointing inwards, versus outwards, on ion conduction, we undertook simulations of the high-K and low-K structures in the presence of high ion concentration (500mM) and high inward-driving membrane potential (-500mV). To trap each structure in the cryo-EM conformation without strongly affecting filter dynamics, weak flat-bottom restraints on S620 H-bonds to the V625-G626 and G626-F627 linkages were applied (not acting unless the conformation attempts to change; see Methods). The filter is seen to conduct ions with the V625 carbonyl oxygens pointing inwards (Fig **1F**; two independent 500ns simulations shown in Supplementary Fig **S6)**. Furthermore, all but one of the seven conduction events observed (Supplementary Fig **S6c**) showed a soft knock-on mechanism with a water molecule separating K^+^ ions during the translocation of ions between S1-S2-S3-S4 K^+^ ion binding sites. As expected, the filter cannot conduct ions when the V625 carbonyl oxygens point outwards (Fig **1F**; two independent 500ns simulations shown in Supplementary Fig **S6**)

In summary, our data, as well as the previous studies of HERG, show a cylindrical shaped selectivity filter in the presence of 300 mM K^+^. Conversely, in the presence of 3 mM K^+^, the selectivity filter has a non-canonical shape with flipped V625 carbonyl oxygens and is not able to conduct K^+^ ions.

### Different Hydrogen bond networks stabilize low-K and high-K structures of HERG

There are multiple residues in the vicinity of the selectivity filter that are either unique to HERG (e.g. S620) or conserved only in the *ether-a-go-go* sub family of voltage gated potassium channels (e.g., Y616, F617, F627, N629; Fig **2A**). Our structures reveal a series of hydrogen bond networks (centred on S620) and hydrophobic interactions (involving F627 and F617) that contribute to stabilisation of the high-K and/or low-K structures. A first set of interactions are present in both the high-K and low-K structures: (i) the S620 backbone carbonyl interacts with the backbone amide of V625 to anchor the bottom of the filter; (ii) the Y616 sidechain hydroxyl and backbone amide of N629 anchor the top of the filter to the pore helix (black dashed lines in Fig **2B**) and (iii) an intersubunit hydrophobic interaction between the sidechains of F627 and F617 cradles the central region of the selectivity filter (Fig **2C**). Together, these interactions form a scaffold that helps to stabilize the structure of the selectivity filter.

**Fig 2.**
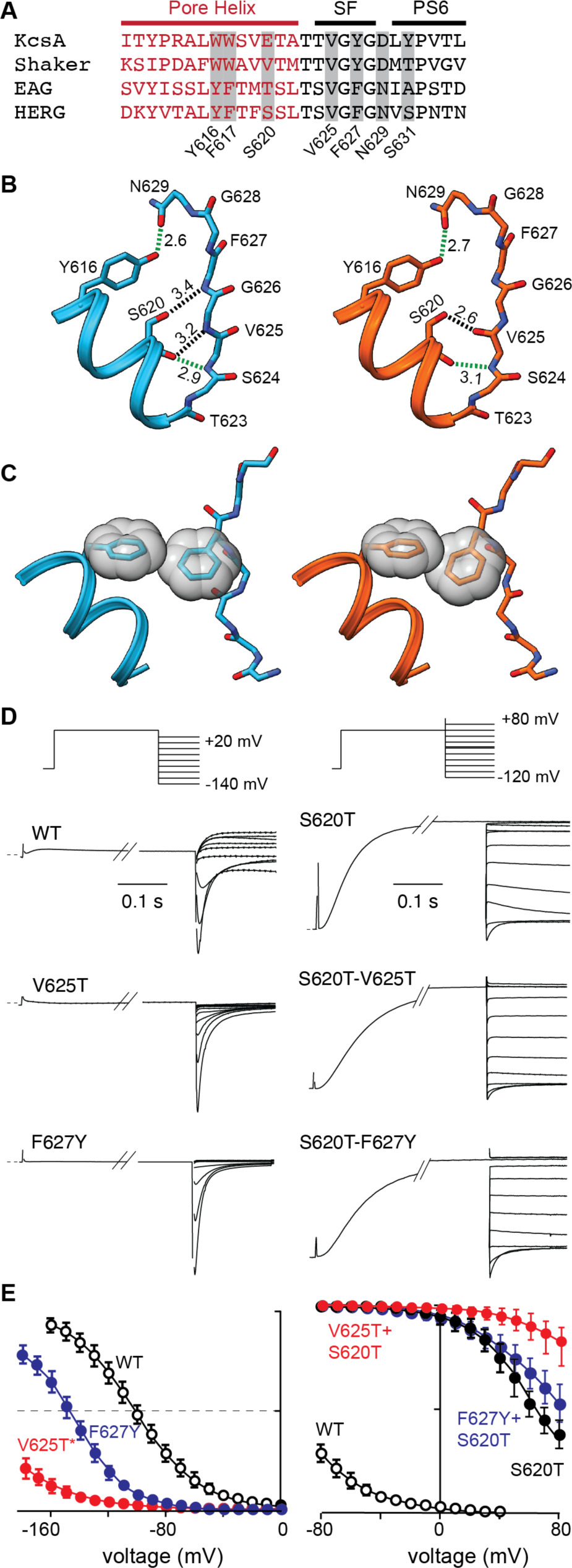
Different hydrogen bond networks stabilize conducting and non-conducting selectivity filters. **A**. Sequence alignment of the pore-helix, selectivity filter and PS6 linker region; residues unique to HERG or present only in the EAG subfamily are highlighted. **B**. Hydrogen bond networks seen in high-K (blue) and low-K (orange) structures. Interactions conserved in both structures (Y616-backbone hydroxyl to N629-sidechain oxygen and S620 backbone carbonyl to V625 backbone amide) are shown in green dashed lines. The S620 sidechain hydroxyl interacts with the G626/F627 backbone amides in the high-K structure but with the flipped V625 backbone carbonyl in the low-K structure. **C.** Intersubunit hydrophobic interactions between the tips of the sidechains of F627 and F617, seen in both high-K (blue) and low-K (orange) structures. **D.** Exemplar families of current traces recorded from V625T and F627Y in WT and S620T backgrounds. In the WT, V625T and F627Y channels pass little current during the initial +40 mV step, as channels are inactivated. Conversely, in S620T, S620T-V625T and S620T-F627Y channels there is a large current at +40mV and these channels only start to inactivate at more positive potentials. NB different voltage range of protocols for S620T channels. **E.** Boltzmann fits to steady-state inactivation curves for mutants shown in panel D. Mutations to V625 and F627 promote inactivation in the WT HERG channels. Conversely, in the presence of S620T, which reduces inactivation, neither V625T nor F627Y enhance inactivation. This is consistent with interactions between S620T-V625T and S620T-F627Y affecting the transition between the conducting and non-conducting states of HERG.

Second, the sidechain of S620 coordinates distinct sets of interactions in the high-K and low-K structures. In the high-K structure, the S620 sidechain interacts with the backbone amides of F627 and G626 whilst in the low-K structure, the S620 sidechain interacts with the flipped V625 backbone carbonyl (Fig **2B**). Notably, mutations to V625 and F627 enhance inactivation when introduced into the WT HERG background, but no longer do so when introduced into the S620T background (Fig **2D**). S620T has a dominant effect on inactivation which indicates that the two mutants are not affecting inactivation via independent mechanisms consistent with V625 and F627 interacting with S620. S620 was first identified as a critical residue for HERG inactivation 25 years ago^8,9^. Here, we have identified how S620 contributes to inactivation gating. Specifically, S620 alternatively stabilizes the canonical SF structure seen in high-K structure through interactions with the amide backbone of G626 and F627 (Fig **2B**), and then stabilizes the flipped V625 carbonyl in the non-canonical SF structure seen in low-K structure (Fig **2B**).

### Effect of K^+^ ions on selectivity filter structure

There are differences in the density along the ion conduction pathway between the high-K and low-K structures when determined with C1 symmetry (Fig **3A**). To explore how changes in K^+^ ion occupancy could affect selectivity filter structure we turned to molecular dynamics (MD) simulations. First, WT channels were embedded in palmitoyloleoyl-phosphatidylcholine lipid bilayers with 1, 2, or 3 ions constrained in the five K^+^ ion binding sites (denoted S0, S1, S2, S3, S4), allowing sampling of the different conformations of the filter (Supplementary Fig **S7**). For simulations with single or multiple ions present, the carbonyl oxygens for the respective ion binding sites tended to point inwards (see Supplementary Fig **S7**). The exception was V625, which contributes to coordination of K^+^ ions in S2 or S3 sites, where some fraction of the carbonyl oxygens were observed to be directed away from the central axis. For example, for S0/S2/S4 occupancy, at least one of the four V625 carbonyl oxygens flips out 84±6% of the time (Fig **3B**). There were no single or multiple ion occupancies that include a K^+^ ion in S2 or S3 where all four V625 carbonyl oxygens pointed towards the axis of symmetry more than 50% of the time. This contrasts with other K^+^ channels where a K^+^ ion held in S2 or S3 is sufficient to stabilize the selectivity filter valine carbonyl oxygens pointing inwards^15,30^. A second notable feature is that the G628 carbonyl oxygens are not constrained to point inwards for any ion configuration (Fig **3B**, Supplementary Fig **S7**) and when ions are not held in S0 or S1, there is a propensity for the F627 carbonyl oxygens to rotate away from the central axis. We also noted during the simulations of ion conduction (Fig **1F**, Supplementary Fig **S6**) that there was a tendency for F627 carbonyl oxygens to rotate away from the pore when ions were not present in S0/S1.These data are consistent with observations from electrophysiology experiments that K^+^ exit from the upper end of the filter represents the first step in the transition from the open to inactivated states of HERG^31^. Lastly, when there are no ions in the filter then the carbonyl oxygens for all residues, apart from S624, are highly dynamic (Supp Fig **S7)**.

**Fig 3:**
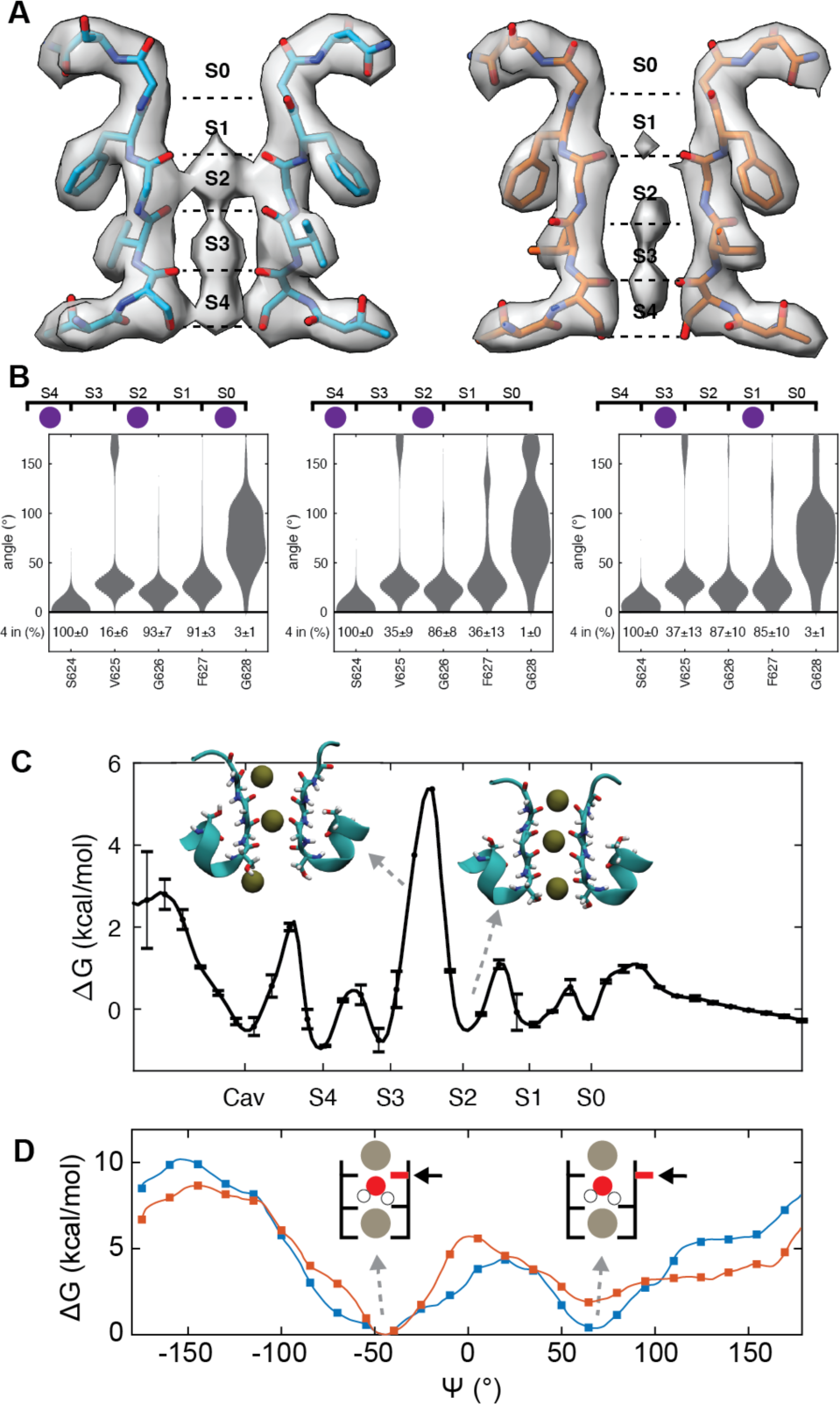
V625 carbonyl oxygens can flip when K^+^ ions are present in the filter. **A.** Structures of high-K (left) and low-K (right) selectivity filters determined with C1 symmetry. The labels S0-S4 highlight canonical K^+^ ion binding sites. **B.** Distribution of carbonyl angles, 8 (0° indicates pointing towards center of the pore) from MD simulations with ions held in the configurations shown. The proportion of time all 4 carbonyl oxygens were pointing inwards (defined as <70° deviation from 0°) is shown at the bottom. Carbonyl angle distributions for single ion configurations are shown in Supplementary Fig **S7**. **C.** Free energy profile for ion binding in the filter from REST2 simulations. There is a large free energy barrier separating S3 and S2 with low free energy barriers between S2, S1, S0 and the extracellular space. **D**. Free energy profile derived from umbrella sampling of rotation of the θ, angle of the V625-Gly626 linkage when ions are held in S0/S2/S4 (blue), or S1/S3/Cav (cavity) (orange). The energy minima for V625 carbonyl oxygens pointing towards and away from the central axis are indicated at θ, =-50° and +70°, respectively.

The above simulations relied on having ions constrained in the filter. To investigate the distribution of unconstrained K^+^ ions in the HERG selectivity filter, and achieve a more unbiased sampling of possible selectivity filter conformations, we used replica exchange with solute tempering (REST2) simulations^32^; totalling 16 µs of enhanced-sampling simulation time. There were frequent movements of ions between sites, as well as ions entering and leaving the filter with an average of 1.8±0.1 ions present in the filter. The free energy profile based on the average ion density shows a large free energy barrier separating S3 and S2 (∼6 kcal mol^-1^, Fig **3C**). As expected, the barrier to K^+^ movement is lowest when all four V625 carbonyl oxygens are pointing inwards; dependence of the free energy profile on the number of V625 carbonyl oxygens directed inward are shown in Supplementary Fig **S8**. The low barriers separating S1 and S0 sites from the extracellular space suggests that ions can rapidly leave S1 and S0 sites. This is consistent with the non-optimal coordination angles of the F627 and G628 carbonyl oxygens in the MD simulations (Supplementary Fig **S7**) and as seen in our low-K structure (Fig **1**). In summary, the combination of a high barrier between S2-S3 and ease of exit of ions from the upper filter provides a plausible explanation for how reducing potassium facilitates HERG adopting a non-conducting filter, as well as the low single channel conductance for HERG channels^33,34^.

All the simulations so far suggest an important role for rotation of the V625 carbonyl oxygens. To directly probe the energetics of this rotation, we used umbrella sampling simulations to map out the free energy of rotation of the V625-G626 peptide linkage. With ions present in the canonical conducting configuration, S0/S2/S4 or S1/S3, there are two minima in the umbrella sampling: at λλ′∼ -50° (open state) and λλ′∼ 70° (inactivated state, Fig **3D**). The free energy difference between flipped and non-flipped V625 carbonyl oxygens is <1 kcal mol^-1^ for the S0/S2/S4 configuration and ∼2 kcal mol^-1^ for S1/S3 configuration; which may be compared to a published estimate of ∼9 kcal mol^-1^ for KcsA^15^. Also, there is only a 4 - 5 kcal mol^-1^ barrier (at λλ′∼ 35°) separating the flipped and non-flipped V625 carbonyl conformations in HERG, compared to ∼12 kcal mol^-1^ for KcsA^15^. Thus, rotation of HERG V625 carbonyl oxygens (both from flipped to non-flipped and vice versa) is energetically possible, when K^+^ ions are present in the filter, and can occur rapidly due to the low energy barrier for flipping.

### The transition state between flipped and non-flipped V625 carbonyl oxygens is facilitated by an interaction between the S620 sidechain and Y616 backbone carbonyl

When we amalgamated the constrained ion MD simulations and clustered frames based on the backbone angles of the selectivity filter residues in individual subunits (Supplementary Fig **S9**), there were two major clusters: Cluster 1 (41% of frames) has carbonyl oxygens of S624, V625, G626 and F627 pointing inwards (Fig **4A**: blue, Supplementary Fig **S9**) and Cluster 2 (30% of frames) has the V625 carbonyl oxygens flipped to point outwards (Fig **4A**: orange. Supplementary Fig **S9**). In addition to the expected interactions between the S620 sidechain and F627/G626 backbone amides (Cluster 1) and with the flipped V625 carbonyl oxygen (cluster 2) we observed that the S620 sidechain spends ∼5% (cluster 1) or ∼25% (cluster 2) of the time interacting with the Y616 backbone carbonyl oxygen (Fig **4B**). When the S620 sidechain is bound to the Y616 backbone carbonyl oxygen, the V625-G626 peptide linkage would be free to rotate, thereby facilitating the transition between the conducting and non-conducting states. To explore this hypothesis, we next analyzed 2D free energy surfaces for S620 sidechain interactions with selectivity filter residues and Y616 as a function of the angle of rotation of the V625 carbonyl in the REST2 simulations which allows more complete sampling of all states (Figure 4**C-F**). The four interactions that the S620 sidechain can participate in, occupy wells in the 2D free energy surfaces that are labelled (i) F627/G626 amides when the V625 carbonyl oxygens point inwards, (ii) Y616 when the V625 carbonyl oxygens point inwards (iii) Y616 when the V625 carbonyl oxygens point outwards and (iv) V625 when the V625 carbonyl oxygens point inwards in Fig **4C-F**. When S620 is interacting with the Y616 backbone, the free energy barrier for valine flipping is only ∼3 kcal mol^-1^, whereas the transition is otherwise forbidden (Fig **4F**). Thus, the S620-Y616 interaction acts like a catalyst to lower the free energy barrier of the transition state and enable rapid rotation of the V625 carbonyl. Mutations to Y616 result in channels that do not express in Xenopus oocytes or in the case of Y616L,I cause a significant perturbation to inactivation gating^35,36^ consistent with this residue playing a critical role in HERG inactivation.

**Fig 4.**
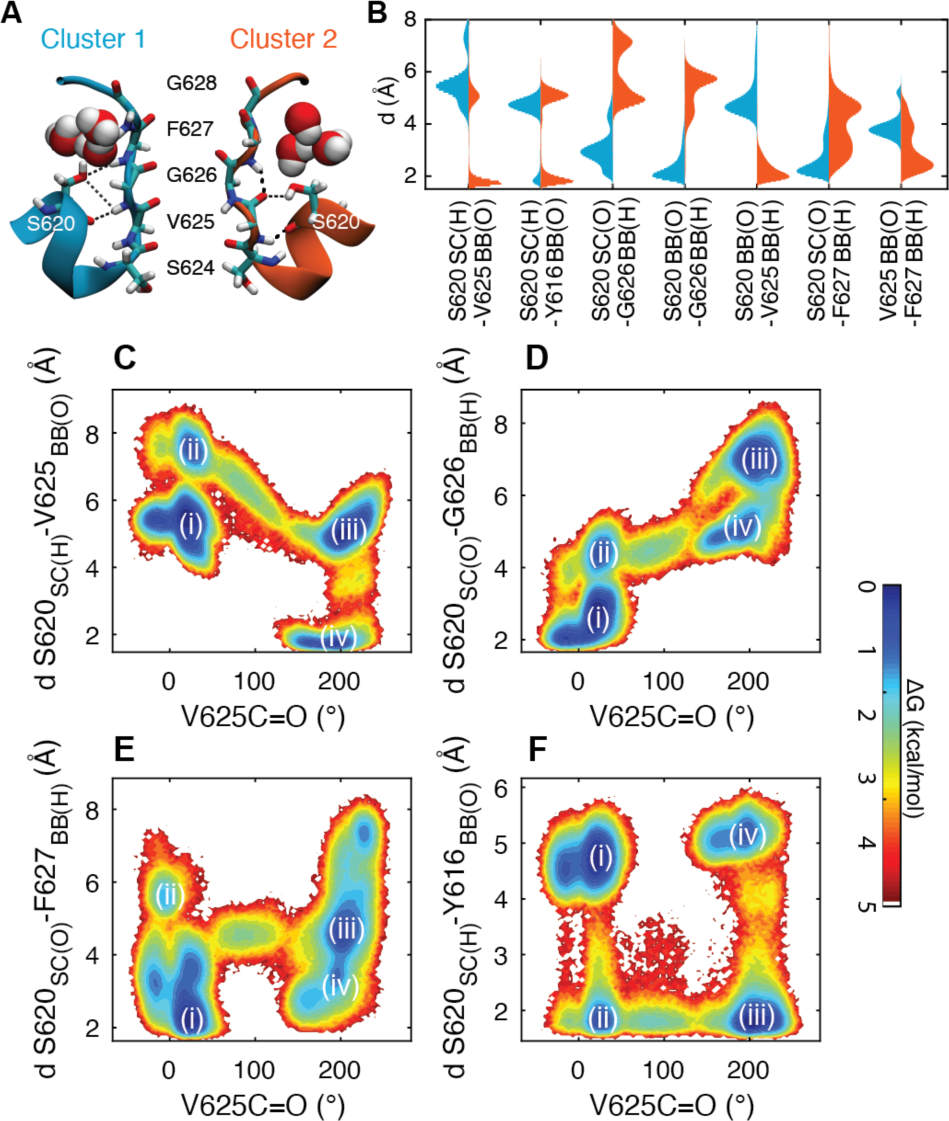
A low energy barrier separates conducting and non-conducting states of the HERG selectivity filters. **A.** Representative structures of the two major clusters observed in constrained ion MD simulations. In Cluster 1 (blue, 41%) all selectivity filter carbonyl oxygens point centrally. Cluster 2 (orange, 30%) has a flipped V625 carbonyl that interacts with S620. **B.** summary of distances between the indicated residue pairs for cluster 1 (blue) and cluster 2 (orange). Note that the S620 sidechain is located ∼2Å from the Y616 backbone carbonyl ∼5% of the time in cluster 1 and ∼25% of the time in cluster 2. **C-F.** Two-dimensional free energy surfaces obtained from REST2 simulations for S620 side chain interactions with **C.** V625 carbonyl, **D.** G626 amide, **E.** F627 amide and **F.** Y616 carbonyl, plotted as a function of rotation of the V625 carbonyl angle (θ). Low free energy states are indicated for S620 interacting with (i) F627/G626, (ii) S620 interacting with Y616 backbone when V625 C=O points inwards, (iii) S620 interacting with Y616 backbone when V625 C=O points outwards, and (iv) S620 interacting with flipped V625 C=O. Very similar results were obtained when we analyzed the amalgamated constrained ion simulations (see Supplementary Fig S10).

### Comparison between WT and S631A HERG

In addition to a structure of WT HERG, Wang and Mackinnon determined the structure of the S631A mutant, which stabilizes occupancy of the open state at 0 mV^37^ In **Fig 5**, we show the selectivity filters of our low-K and high-K WT structures and compare to the previously determined high-K S631A structure (PDB: 5VA3). The three structures have been aligned based on the pore helix region. In addition to rotation of the F627 residue between S631A and WT high-K (as noted previously^22^) the WT HERG filter is displaced upwards relative to the filter in S631A. The higher displacement of the WT HERG filter raises the level of the V625 carbonyl oxygens to be closer to the level of S620 sidechain. Thus, we suggest that the higher filter in the WT high-K structure is better placed to allow rotation about the V625-G626 peptide linkage and so can be considered primed for inactivation as soon as K^+^ ions leave the filter. Conversely, we suggest that in the S631A channel, the lower filter would promote interaction between S620 and the amide backbones of G626 and F627 to stabilize the conductive filter. This model needs to be tested by investigating how mutations to residues that are known to affect inactivation gating, both within the filter region as well as in the pore helix, PS6 domain and the extracellular turret region affect the structure of the selectivity filter.

**Fig 5:**
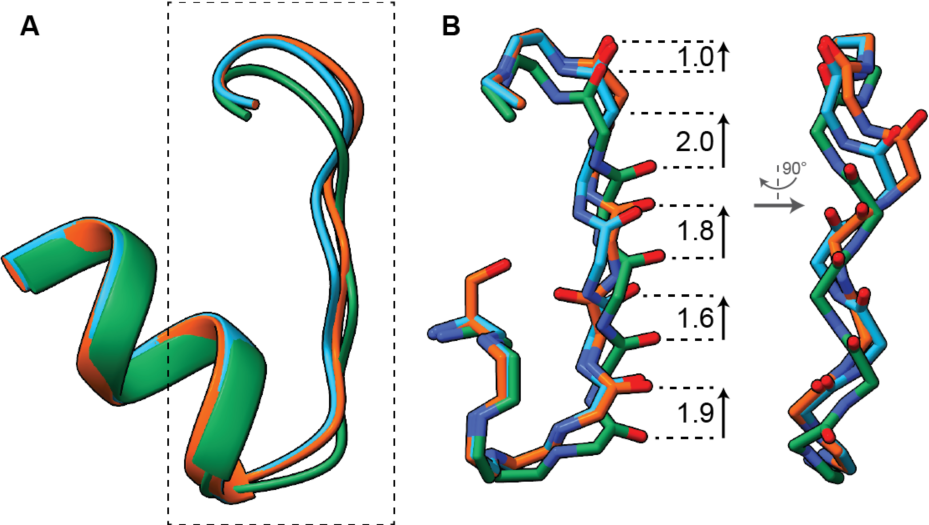
Comparison between selectivity filter structures in WT high-K, WT low-K and S631A HERG. **A.** The three structures (orange: WT Low-K, blue: WT High-K, green: S631A, 5va3.pdb) were aligned on the pore helix regions. The S631A filter (green) is displaced downwards relative to the WT structures. **B.** Highlight of the boxed region from A, showing the relationship between the backbone atoms in the selectivity filter. Distances on left side indicate vertical displacement of backbone Cα atoms for S631A and WT low-K

## Discussion

From the cryo-EM and MD studies, we propose a model for the K^+^-dependent transition between conducting and non-conducting states of HERG that is summarized in **Fig 6**. The conducting state is characterized by the S620 sidechain interacting with the amide backbones of F627 and G626 (**Fig 6a**). Transition from conducting to non-conducting is initiated by K^+^ ions leaving the upper filter (**Fig 6b**). The transition state (**Fig 6c,d**) is characterized by the S620 sidechain interacting with the Y616 backbone, which leaves the Val-Gly peptide bond free to rotate. The inactivated state is stabilized by the flipped V625 carbonyl binding to the S620 sidechain. The high barrier between S2-S3, which slows ions moving upwards, coupled with the ease of exit of ions from the upper filter to the extracellular solution, means that ions may leave and not be replenished quickly, leaving the upper filter free of ions for a period, which facilitates transition to a non-conducting state. Importantly, our model explains the critical role for S620^8,9^, which is validated by the observation that this residue is unique to HERG and that the homologous HEAG channels, where the corresponding residue is a threonine, do not inactivate ^8^. We have also identified a transition state, formed by the S620 sidechain interacting with the Y616 backbone carbonyl oxygen that leaves the V625-G626 peptide linkage free to rotate between the conducting and non-conducting states. Our model is also supported by an extensive literature showing that mutations to the residues involved in the interactions described above (Y616, S620, V625, G626, F627) affect inactivation gating in HERG^8,24,35,36,38,39^.

**Figure 6:**
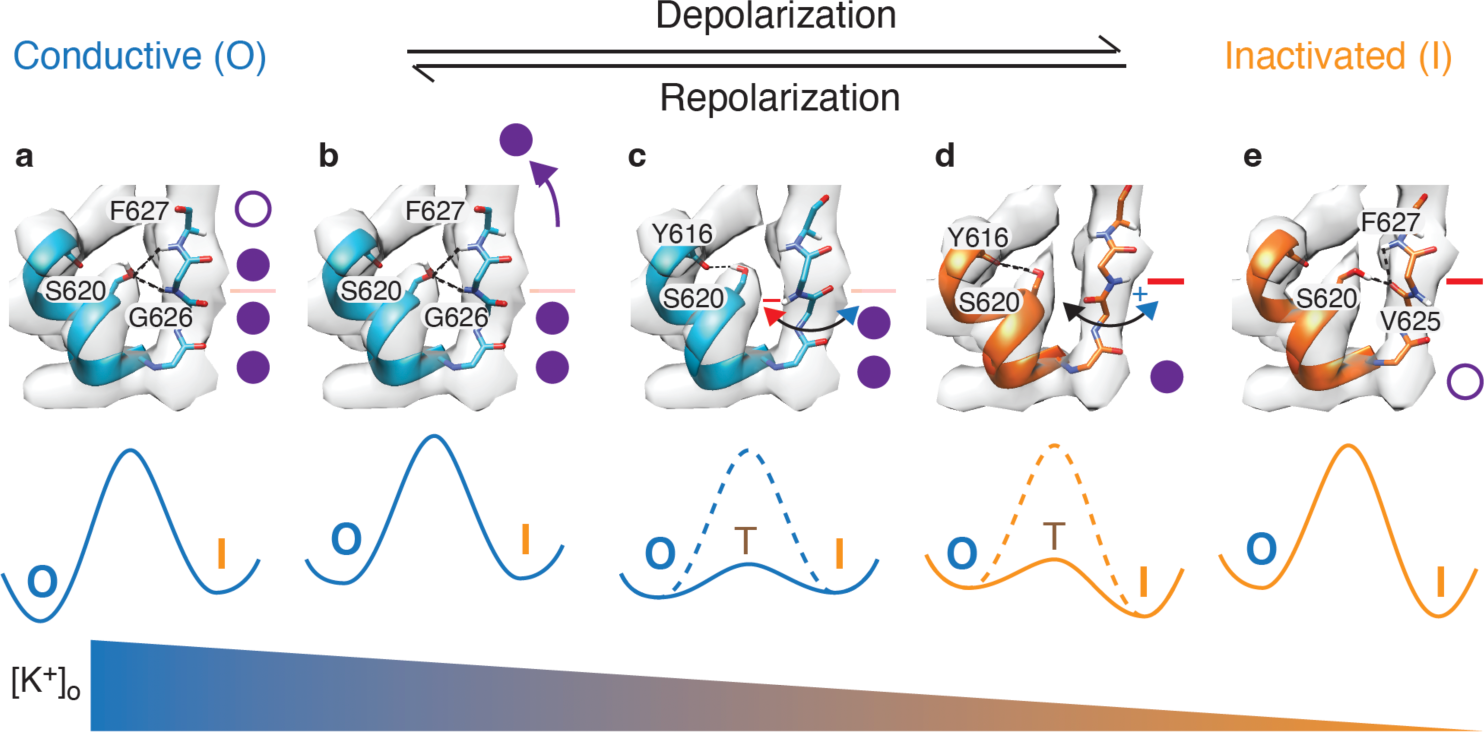
Proposed model for transition from conducting to non-conducting selectivity filter. Cryo-EM maps of the selectivity filter obtained from the high-K structure (panels **a-c**) and low-K structure (panels **d-e**) with superimposed protein structures obtained from snapshots of the MDFF fits to the high-K (blue) or low-K (orange) cryo-EM maps. Stylized free energy diagrams for the transition from the conductive (O) to inactivated (I) state, via a transition state (T) are shown below each panel. Interactions stabilizing each state are highlighted as dashed lines in each panel (**a,b**: S620 – F627/G626, **c,d**: S620 – Y616; **e**: S620-V625). Inactivation is initiated by K^+^ ions leaving the upper filter (**b**). The energy barrier for movement of ions between S3 and S2 (pink line, panels **a,b**) increases when the V625 carbonyl oxygens are flipped (red line in panels **d,e**) The gradient for [K^+^]o indicates that the conductive state is favored at high [K^+^]o and the inactivated state is favored at low [K^+^]o. A movie depicting the transition from the conductive to inactivated state is shown in Supplementary Movie 1.

Based on crystallography studies of KcsA in high (150-300 mM) and low (3 mM) potassium, inactivation was proposed to involve constriction of the central glycine residue^10,14^, rotation of V76 carbonyl oxygens away from the central axis^10,11,17,40,41^), rearrangement of water-mediated hydrogen bond networks, and decrease in the number of water molecules behind the selectivity fitler^11,14,17^. In HERG, there is more rotation of the V625 carbonyl oxygens but a less prominent constriction at the level of G626 (for both the carbonyl oxygens and backbone Cα), compared to KcsA (**Fig. 7**). The interaction between the flipped V625 carbonyl oxygens and the S620 sidechain hydroxyl in HERG may also contribute to the “non-conducting” conformation of HERG being longer lived compared to the flipped V76 carbonyl oxygens in KcsA and hence why the flickering state seen in KcsA is not as prominent in HERG^33,34^. It is not possible to observe water molecules at the resolution of our cryo-EM structures, but our MD simulations suggest that there is only a modest rearrangement with no net change in the number of water molecules behind the filter during the transition between conducting and non-conducting states (Supplementary Fig 11). The no net change in water molecules may contribute to kinetics of HERG inactivation being faster than that observed for KcsA where the rate of inactivation is determined by the diffusion limited rate of water binding^17^.

**Figure 7:**
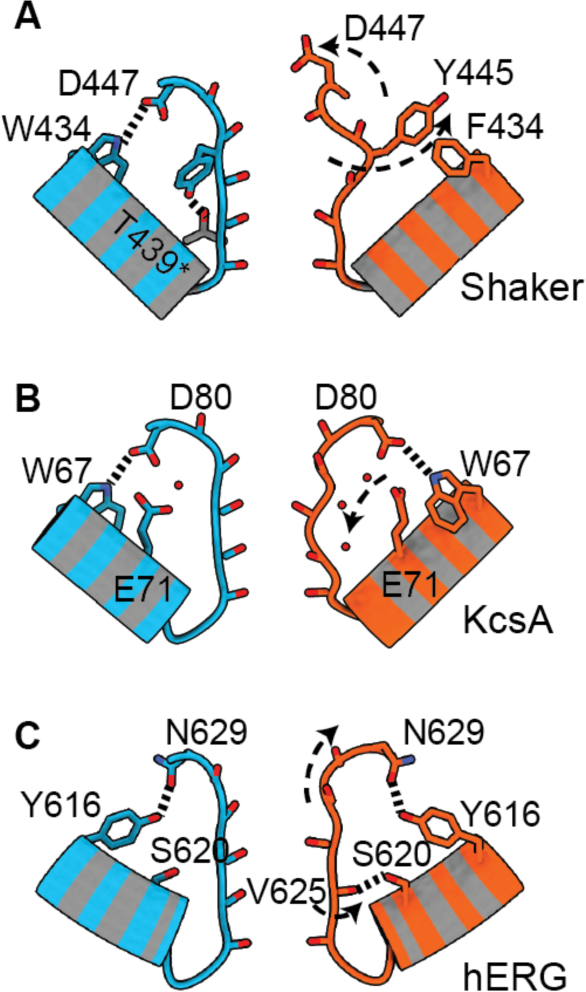
Different channels adopt different non-conducting selectivity filter structures. **A.** In Shaker channels there is dilatation of the central glycine and tyrosine carbonyl oxygens but little change in the lower filter^18,19^. In KcsA, the non-conductive state is characterized by constriction of the central glycine carbonyl oxygens, an incomplete rotation of the valine carbonyl oxygens away from the central axis ^10^ and changes in water occupancy ^17^. The non-conductive state in HERG is more like KcsA than Shaker but involves a complete rotation of the V625 carbonyl oxygens, stabilized by an interaction with the S620 sidechain and no change in the number of water molecules.

S620 in HERG is equivalent to E71 in KcsA and both play critical roles in inactivation, albeit in different ways that reflect the distinctive physiochemical properties of the respective sidechains. In two previous MD studies of HERG, it was proposed that S620 might interact with N629 directly^23,42^ or via a water-mediated interaction^23^, analogous to the E71-D80 interaction seen in WT KcsA^10,11,17^. Given the shorter S620 sidechain (compared to E71) an interaction between S620 and N629 results in a buckling of the selectivity filter in HERG^23,42^. Rather than interaction with S620, our structures show an interaction between N629 and Y616 that is preserved in both the high and low-K structures (**Fig 3**) and so could serve to scaffold the upper part of the filter to the pore-helix in both conducting and non-conducting states. We also note that there is an interaction between the homologous residues in KcsA (D80 and W67)^10,11,14^ that can be seen in both conducting and non-conducting filters.

Comparison between HERG and the recent structures of Shaker, and the homologous Kv1.3, are more tenuous. In Shaker, the V438 sidechain (equivalent to S620 in HERG) cannot form hydrogen bonds and the putative inactivated state appears to have a distinct structure with dilatation of the upper filter^18–21^. The published structures for the “inactivated” state of Shaker, however, are based on mutant structures, rather than changes in K^+^ concentration. In a recent MD simulation study of HERG, Pettini and colleagues showed that mutants that promoted inactivation led to widening of the extracellular entrance of the channel, but less prominent than what was seen in Shaker^25^. In our cryo-EM structures we observed a small dilatation of the upper filter, in both the high-K and low-K structures (**Supplementary Fig S5**). Both our data and the MD simulations from Pettini et al.^25^ are consistent with the conducting state of the filter being sensitive to loss of K^+^ from the extracellular side, which has been shown to represent the first step in the transition from the open to inactivates state^31^.

The other channel for which high-K and low-K structures are available is the heterodimeric K2P.1 (Trek1) channel^12^. These channels undergo asymmetric changes with pinching of one of the two selectivity filter sequences but dilatation of the second selectivity filter sequence, further supporting the hypothesis that all K^+^ channels adopt a similar conductive conformation, but each adopt different non-conducting filter configurations. It has been suggested that the non-conducting selectivity filter of HERG may involve asymmetric changes^24^. In our MD studies, the flipping of the V625 carbonyl oxygens occurred in individual subunits, which resulted in some asymmetric changes, The asymmetry that we observed, however, was small, including in the extensive REST2 simulations which sampled broad-ranging configurations of the selectivity filter (Supplementary Fig **S12**). This is consistent with the lack of significant asymmetry in the low-K structure determined by cryo-EM.

In summary, the structure of the conducting filters for KcsA, Shaker and HERG are comparable to each other, but the non-conducting filters are different to each other. These subtle differences can explain the uniquely rapid inactivation kinetics of HERG channels and paves the way for future investigations into understanding how clinically occurring mutations in the vicinity of the selectivity filter, as well as movement of the activation gate, alter the distribution between conducting and non-conducting states of HERG.

### Key resources table

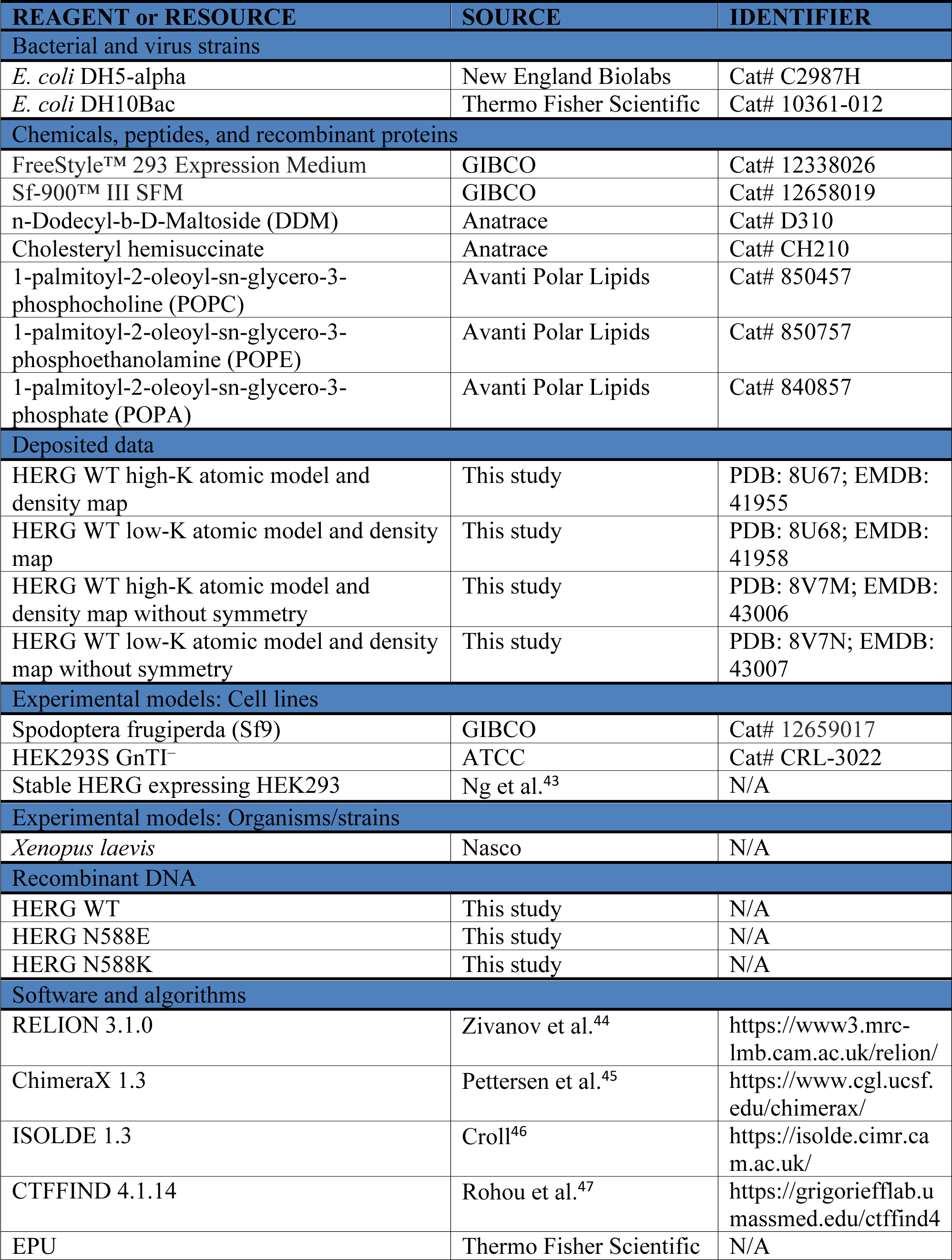
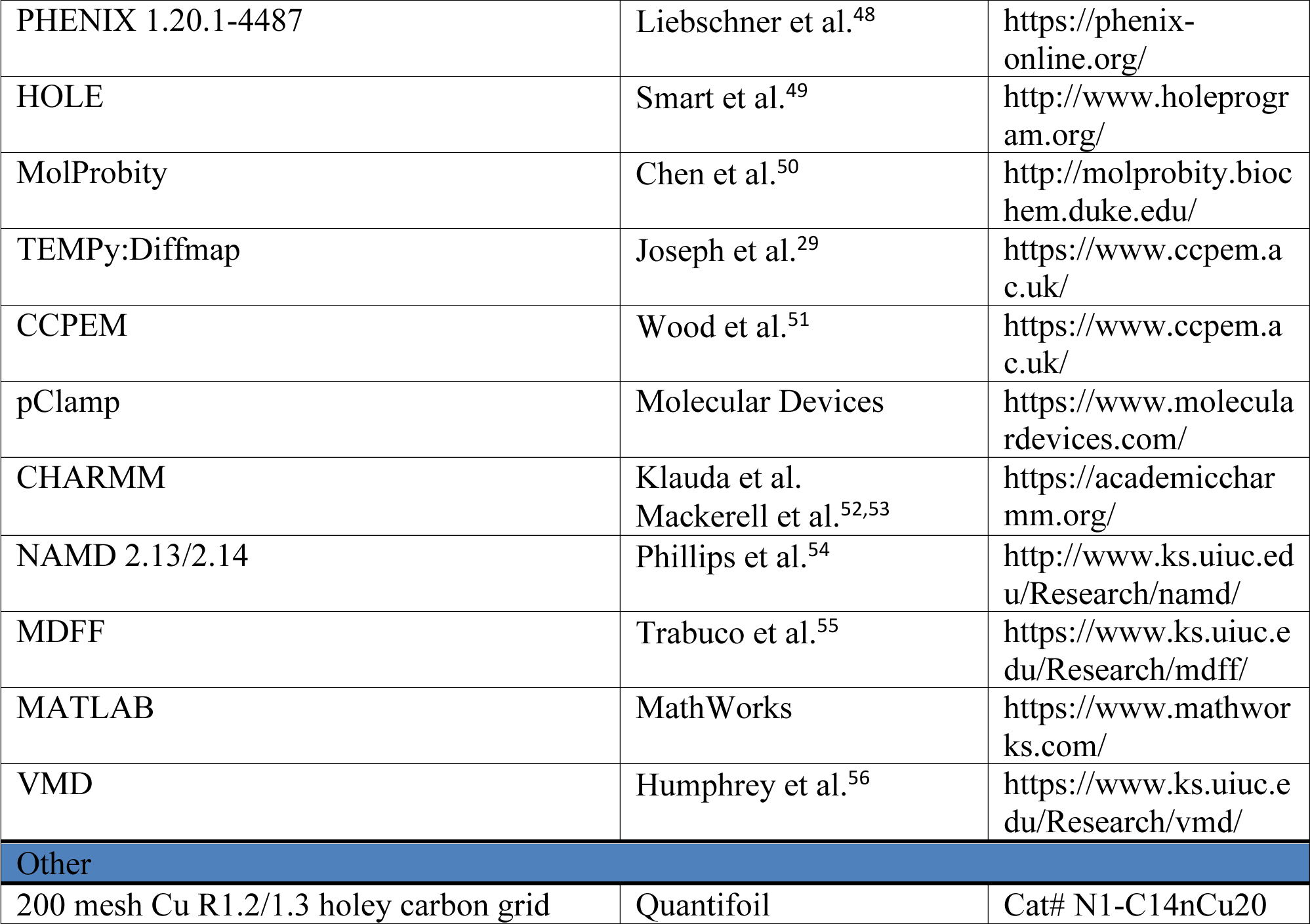

| REAGENT or RESOURCE | SOURCE | IDENTIFIER |
| --- | --- | --- |
| Bacterial and virus strains |  |  |
| *E. coli* DH5-α | New England Biolabs | Cat# C2987H |
| *E. coli* DH10Bac | Thermo Fisher Scientific | Cat# 10361-012 |
| Chemicals, peptides, and recombinant proteins |  |  |
| FreeStyle™ 293 Expression Medium | GIBCO | Cat# 12338026 |
| Sf-900™ III SFM | GIBCO | Cat# 12658019 |
| n-Dodecyl-b-D-Maltoside (DDM) | Anatrace | Cat# D310 |
| Cholesteryl hemisuccinate | Anatrace | Cat# CH210 |
| 1-palmitoyl-2-oleoyl-sn-glycero-3-phosphocholine (POPC) | Avanti Polar Lipids | Cat# 850457 |
| 1-palmitoyl-2-oleoyl-sn-glycero-3-phosphoethanolamine (POPE) | Avanti Polar Lipids | Cat# 850757 |
| 1-palmitoyl-2-oleoyl-sn-glycero-3-phosphate (POPA) | Avanti Polar Lipids | Cat# 840857 |
| Deposited data |  |  |
| HERG WT high-K atomic model and density map | This study | PDB: 8U67; EMDB: 41955 |
| HERG WT low-K atomic model and density map | This study | PDB: 8U68; EMDB: 41958 |
| HERG WT high-K atomic model and density map without symmetry | This study | PDB: 8V7M; EMDB: 43006 |
| HERG WT low-K atomic model and density map without symmetry | This study | PDB: 8V7N; EMDB: 43007 |
| Experimental models: Cell lines |  |  |
| Spodoptera frugiperda (Sf9) | GIBCO | Cat# 12659017 |
| HEK293S GnTI<sup>-</sup> | ATCC | Cat# CRL-3022 |
| Stable HERG expressing HEK293 | Ng et al.<sup>43</sup> | N/A |
| Experimental models: Organisms/strains |  |  |
| *Xenopus laevis* | Nasco | N/A |
| Recombinant DNA |  |  |
| HERG WT | This study | N/A |
| HERG N588E | This study | N/A |
| HERG N588K | This study | N/A |
| Software and algorithms |  |  |
| RELION 3.1.0 | Zivanov et al.<sup>44</sup> | https://www3.mrc-lmb.cam.ac.uk/relion/ |
| ChimeraX 1.3 | Pettersen et al.<sup>45</sup> | https://www.cgl.ucsf.edu/chimerax/ |
| ISOLDE 1.3 | Croll<sup>46</sup> | https://isolde.cimr.cam.ac.uk/ |
| CTFFIND 4.1.14 | Rohou et al.<sup>47</sup> | https://grigoriefflab.umassmed.edu/ctffind4 |
| EPU | Thermo Fisher Scientific | N/A |
| PHENIX 1.20.1-4487 | Liebschner et al.<sup>48</sup> | https://phenix-online.org/ |
| HOLE | Smart et al.<sup>49</sup> | http://www.holeprogram.org/ |
| MolProbity | Chen et al.<sup>50</sup> | http://molprobity.biochem.duke.edu/ |
| TEMPy:Diffmap | Joseph et al.<sup>29</sup> | https://www.ccpem.ac.uk/ |
| CCPEM | Wood et al.<sup>51</sup> | https://www.ccpem.ac.uk/ |
| pClamp | Molecular Devices | https://www.moleculardevices.com/ |
| CHARMM | Klauda et al.<br>Mackerell et al.<sup>52,53</sup> | https://academiccharmm.org/ |
| NAMD 2.13/2.14 | Phillips et al.<sup>54</sup> | http://www.ks.uiuc.edu/Research/namd/ |
| MDFF | Trabuco et al.<sup>55</sup> | https://www.ks.uiuc.edu/Research/mdff/ |
| MATLAB | MathWorks | https://www.mathworks.com/ |
| VMD | Humphrey et al.<sup>56</sup> | https://www.ks.uiuc.edu/Research/vmd/ |
| Other |  |  |
| 200 mesh Cu R1.2/1.3 holey carbon grid | Quantifoil | Cat# N1-C14nCu20 |

## RESOURCE AVAILABILITY

### Lead contact

Further information and requests for resources and reagents should be directed to and will be fulfilled by the lead contact, Jamie I. Vandenberg.

### Materials availability

All unique/stable reagents generated in this study are available from the Lead Contact with a completed Materials Transfer Agreement.

### Data and code availability

Cryo-EM maps have been deposited at Electron Microscopy Data Bank (EMDB) and are publicly available as of the date of publication. Accession numbers are listed in the key resources table.

Coordinates of atomic models have been deposited at Protein Data Bank (PDB) and are publicly available as of the date of publication. Accession numbers are listed in the key resources table.

Any additional information required to reanalyze the data reported in this work paper is available from the lead contact upon request.

## METHOD DETAILS

### Electrophysiology studies

*Xenopus oocyte recordings:* WT HERG cDNA was a gift from Gail Robertson (University of Wisconsin). Mutants were generated by Mutagenex Inc. (Piscataway, NJ). cRNA was synthesized using the mMessage mMachine kit (Ambion) according to the manufacturers’ protocols. Oocytes from *Xenopus laevis* frogs (2-6 years of age; purchased from Nasco, Fort Atkinson, WI, USA) were prepared as previously described^57^. The Garvan/St Vincent’s Animal Ethics Committee approved all experiments. All experiments were undertaken at room temperature (21-22°C). Perfusion solutions contained (*x* mM KCl, 100-*x* mM NaCl, 1.8 mM CaCl_2_, 1 mM MgCl_2_, 5 mM HEPES, pH adjusted to 7.5 with NaOH) where *x* was 3 or 100. Glass microelectrodes had tip resistances of 0.2 – 0.7 MΩ when filled with 3M KCl. Data analysis was performed using pClamp software (Version 10.6, Molecular Devices) and Excel software (Microsoft Corporation, Seattle, WA). All data are shown as mean ± S.E.M.

*Steady-state inactivation* was measured from a two-step voltage protocol, as previously described^57^. The voltage range for measurement of inactivation in Xenopus oocytes extended to -200 mV for mutants with enhanced inactivation and up to +100 mV for mutants with reduced inactivation. For HEK293 cells inactivation was measured over a voltage range from -150 mV to +20 mV. Peak tail currents were obtained by fitting an exponential function to the decay phase of the tail currents and extrapolating back to the start of the voltage step, i.e., to account for channel deactivation^2,4^. Normalized conductance values were then fitted with a Boltzmann function:

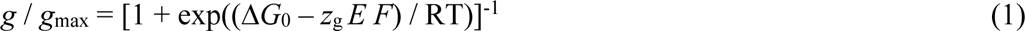

where Δ*G*_0_ is the free energy difference between the open and inactivated states at 0 mV. *z*_g_ is the charge transferred during inactivation, *E* is the electric field strength, *F* is Faraday’s constant, *R* is the Universal Gas constant and *T* is temperature.

### Protein expression and purification

The cryo-EM construct used for the study of Human KCNH2 (HERG) was based on the Ts construct developed by Wang and Mackinnon^22^. In brief cytoplasmic loops 141-380 and 871-1005 were removed using quikchange PCR methods. A TEV protease site (ENLYFQG) was inserted between the gene and a GFP epitope followed by a double strep tag (AWSHPQFEK) with a linker (GGGS)_2_ (GGSA) between the first and second repeat. The construct was cloned using EcoR1 and Xba1 into PEG vector^58^.

The PEG-HERG construct was used to generate baculovirus as described in^58^. Cells were cultured and purifications were performed with changes as described^22^. Briefly, HEK 293 GnTI were induced with baculovirus when reaching a density of 2.5-3.5 × 10^6^ cells/ml. After 24 hours from induction the temperature was lowered to 30°C and a final concentration of 10 mM sodium butyrate was added for a further 24 hours where cells were harvested.

Cells were lysed in 30 mM KCl and 20 mM HEPES pH 7.4 and spun down at 40,000 rpm for 90 mins. The insoluble fraction was collected and resuspended in 300 mM KCl, 20 mM HEPES pH 7.4, 1% DDM n-Dodecyl-b-D-maltoside (DDM), 0.2% Cholesteryl hemisuccinate (CHS) and 5mM DTT, samples were incubated at 4°C for one hour under gently agitation. The soluble fraction was collected and loaded onto strep-tactin superflow high-capacity resin (IBA) and allowed to bind at a flow rate of 1 ml/min at 4 degrees. Resin was washed (300 mM KCl, 20 mM HEPES pH 7.4, 0.1% DDM, 0.02% Cholesteryl hemisuccinate (CHS), 5 mM DTT and 0.1 mg/ml phospholipids POPC, POPE and POPA in a ratio of 5:5:1) with 5 column volumes or until UV absorption returned to baseline. The sample was eluted with an addition of 5 mM Desthiobiotin to the wash buffer. For WT HERG, GPF was cleaved with TEV protease overnight at 4°C under gentle agitation. The Tetramer peak was collecting using a Superose 6 using either 300 mM KCl or 3 mM KCl and 297 mM NaCl, 20 mM HEPES pH 7.4, 10 mM DTT, 0.025% DDM, 0.005% CHS and 0.025 mg/ml of phospholipids POPC, POPE and POPA in a ratio of 5:5:1. Protein was concentrated to 7.5 mg/ml.

### Cryo-EM grid preparation and data collection

Quantifoil® 200 Cu mesh R1.2/1.3 holey Cu-carbon grids were plasma cleaned for 60 s in low pressure gas (0.5 mbar, 80% argon 20% oxygen mixture) using a Diener plasma cleaner. Protein sample (3.5 µl; 7.5 – 8.5 mg/ml) was applied to the carbon coated side of the grid. The grid was blotted for 10 s with blot force 10 and vitrified in liquid ethane using a Vitrobot Mark IV (FEI) equilibrated to 4 °C and 100% humidity.

For all datasets, grids were imaged on a Titan Krios operating at 300 keV equipped with a Gatan K2 detector. Images were collected in electron counting mode at a nominal microscope magnification of 130kx (1.05 Å/pixel) with 1 e^-^/A^2^ per frame with a total dose of 50 e^-^/A^2^ or 60 e^-^/A^2^ and nominal defocus range from -0.5 to -2.5 µm.

### Cryo-EM data processing

Data was processed using RELION-3.1.1. Motion correction was done using RELION^44^ and defocus values were estimated using CTFFIND4^47^. Auto-picking was first performed on a subset of micrographs for each datasets using a Laplacian of Gaussian filter to generate templates for templated-based auto-picking of the whole dataset. Particles were extracted from micrographs with a box size of 300 pixels and binned to 64 pixels for all datasets The binned particles were subjected to multiple rounds of 2D classification. Good classes were manually selected and reextracted without binning. An initial model was generated without imposing symmetry. The extracted particles were initially subjected to one round of 3D classification with 6 classes and no symmetry imposed. No asymmetry was observed in the initial model or any of the classes (see Supplementary Fig **S1**). Further 3D classification was therefore performed with C4 symmetry imposed and classes which clearly resemble HERG were selected. Final 3D auto-refinement followed by Bayesian polishing and CTF refinement were performed in RELION.

Post-processing and resolution estimates were performed with a soft mask including only the transmembrane domains. Pixel size was calibrated against the published structure (EMD-8650) by maximizing the cross-correlation between the two maps. Calibrated pixel size of the final map was 1.05 Å/pixel for all datasets.

### Model building and refinement

Model was built and refined using ISOLDE^46^ and PHENIX^48^. The HERG structure (PDB: 5VA1) was used as starting model. Only the transmembrane domains (residues 398-709) were modelled due to the poorer local resolution for the cytoplasmic domains. MolProbity^59^ was used to validate the geometries of the refined models. Difference maps were calculated using TEMPy:Diffmap^29^. The WT high-K and WT low-K maps were first low pass filtered to 3.3Å, then matched by amplitude scaling in resolution shells and the difference map generated as fractional differences with respect to the globally scaled map values. All images were rendered using ChimeraX-1.4^45^. Internal cavity was calculated using HOLLOW ^60^ with 1.4 Å radius probe and grid spacing of 0.2 Å.

### Molecular dynamics

All systems were built with CHARMM and simulated with NAMD2.13 or NAMD2.14^54^. The CHARMM36 lipid^52^ and CHARMM22 protein force fields^61^ with CMAP corrections^53^ were used with modified K^+^-carbonyl interaction parameters (depth 0.102 kcal/mol and position 3.64 Å of minimum) to achieve a small experimental preferential solvation of K^+^ in N-methyl-acetamide over water^62^.. The NPT ensemble was maintained using the Langevin piston Nose-Hoover method^63,64^ for pressure and Langevin dynamics to maintain a temperature of 303 K. Bonds to hydrogen atoms were maintained with the RATTLE algorithm^65^ and electrostatic interactions calculated with Particle Mesh Ewald^66^ with a grid spacing of 1.0 Å and 6th order B-spline mesh interpolation with a neighbor list distance of 15 Å and a real space cut-off of 12 Å with energy switch distance of 10 Å.

### Molecular Dynamics Flexible Fitting

To best mimic the experimental system for Molecular dynamics flexible fitting (MDFF)^55^ simulations, detergent micelles were built around proteins using PDB:5VA1. The optimal size of the micelle was determined by simulating 6 different sizes of a simple pure detergent micelle (n-Dodecyl-B-Maltoside Detergent (DDM). The number detergent molecules in contact with the protein plateaus at 400-450 molecules, Supp Fig **S4**, suggesting that this is enough micelle molecules for embedding HERG^67^. WT HERG, N588K_HERG and N588E_HERG were embedded in micelles containing 404 molecules, consisting of detergent (DDM (300)), lipid (POPE (20), POPC (20), POPA (4)) and cholesterol derivative (cholesteryl hemisuccinate (60)), and surrounded with explicit TIP3P water molecules^68^. Micelles were surrounded with 110 853 water molecules and either 660 K^+^ ions and 600 Cl^−^ ions (300 mM KCl) or 70 K^+^ ions and 6 Cl^−^ ions (3mM KCl) respectively.

MDFF^55^ was used to refine the cryo-EM structures in the presence of a potential energy function based on the cryo-EM density map Φ(**r**), given by *U_EM_*(***R***) = ∑ω_*j*_*V*_*EM*_(**r**_*j*_), where *ω* is the weight for each atom

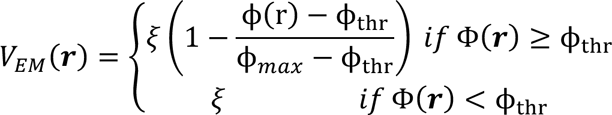

with threshold, Φ_*+,_, to remove noise and scaling factor ξ. Five independent simulations were performed for each map, where ξ was progressively increased from 0 to 5 kcal/mol over the first 50 ns followed by 50 ns of constant ξ=5 kcal/mol. For the WT simulations, the RMSD for the backbone atoms of the pore domain helices and the selectivity filter plateaued after 10 ns and 40 ns, respectively. 50 final structures were generated by energy minimising with 1000 steps of steepest descent every ns with ξ =10 kcal/mol starting after 50 ns simulation. For N588K and N588E HERG, a final structure was saved from an MDFF simulation to use as the starting point for subsequent K^+^ ion permeation US simulations (see below).

### Molecular dynamics simulations of HERG in membranes

#### Simulations of HERG with maintained ion configurations

MD simulations were performed starting with the WT HERG high-K structure. The pore domain (residues 545 to 667) was embedded in a lipid bilayer consisting of palmitoyloleoyl-phosphatidylcholine (POPC) lipids (238 molecules). It was surrounded with 18,771 explicit TIP3P water molecules^68^ and 150 mM KCl (49 K^+^ ions and 53 Cl^−^ ions). During initial equilibration, all heavy atoms in the protein, as well as ions in the selectivity filter, were constrained with harmonic constraints with force constants of 10 kcal/mol/Å^2^. These were slowly released on the protein during 5 ns simulations before production runs.

Libraries of 10 independent simulations each of 50 ns were performed for the WT channel with each of 9 different ion configurations in the selectivity filter. Multi-ion configurations in sites S0/S2/S4, S2/S4 and S1/S3 as well as single ion configurations in sites S0-S4 and an empty filter were constrained with 5 kcal/mol in their sites with flat bottomed potentials (*k* = 10 kcal/mol/Å^2^ and thickness 3.5 Å). Means and distributions were calculated after excluding the first 10 ns of equilibration. Errors bars were calculated as the standard error of means for the 10 different simulations for each ion configuration.

The angle of the selectivity filter carbonyl oxygens, θ, were calculated as the angle between the projection on the *xy*-plane of the vector between the carbonyl C and the carbonyl O and the projection on the *xy*-plane of the vector between the carbonyl C and the center of mass of the selectivity filter. Distributions of carbonyl angles (0° indicates pointing towards center of pore) have been plotted as violin plots, and the proportion of time 0, 1, 2, 3, or 4 carbonyl oxygens were pointing inwards (defined as <70° deviation from 0°) reported.

Distances between key residues were calculated and binned as a 2-D density, ρ, together with the angle of the V625 carbonyl. 2-D free energy maps were calculated from this density by Δ*G* = −*k_B_Tln*(ρ).

Cluster analysis was performed with k-means algorithm^69^ in MATLAB using *k* = 10 clusters and clustering on the backbone <λ and 4¢¢ angles of the selectivity filter (residue 624 to 628). Initial cluster centroids were generated using the k-means++ algorithm. Each point in space was then allocated to its nearest cluster and the cluster centroid recalculated. This was done iteratively until self-consistent. Each subunit was clustered separately to allow for asymmetries. Analysis of carbonyl angles and interaction distances (*d*) was performed based on which cluster the system was in (blue conducting, orange non-conducting). Free energy maps, as function of S620 side chain interactions with V625 carbonyl, G626 amide, F627 amide or Y616 carbonyl plotted against rotation of the V625 carbonyl have been obtained from analysis of all frames of the constrained ion MD simulations of HERG in the membrane. Maps were created from calculation of Δ*G*(*d*, θ) = −*k_B_T* ln ρ(*d*, θ) + *C*, where *ρ* is the probability distribution as a function of the interaction distance *d* and V625 carbonyl orientation θ.

#### Conduction simulations

Simulations starting with the high-K structure were similar to those described above, but included 500mM KCl solution (165 K^+^, 169 Cl^−^ ions and 18,653 water molecules), including 5 K^+^ ions that were initially placed at the centres of sites S0-S4 in the equilibrated structure. The CHARMM36 force field without modifications was used for these simulations. This system was run for 0.5 μs (two independent simulations) with a constant electric field equivalent to an applied membrane potential of -500mV (negative inside) to attempt to observe conduction. During equilibration, heavy atom harmonic position restraints on the selectivity filter and resident ions were applied with a force constant of 5 kcal/mol/Å^2^, relaxed to zero slowly over 10 ns. To maintain this high-K cryo-EM-like structure, flat-bottom restraints were applied to loosely maintain H bonds identified in the cryo-EM structure. This required a weak flat-bottom restraint (force constant 5 kcal/mol/Å^2^ applied when 3.2 Å was exceeded) acting on the distance between the S620 side chain hydroxyl O, and both the G626 and F627 backbone amide H atoms for each subunit, for the conducting structure. This restraint is designed to not be felt unless the H-bond is attempting to break, as would occur during a structural isomerization of the backbone linkage. In addition, a flat-bottom harmonic restraint was applied to the intracellular gate. Specifically, the 6 Cα-Cα distances connecting adjacent carbon atoms of the same residue were weakly restrained with flat-bottom distance restraints, for the four Cα atoms (each) in residues F656, G657 and N658. The 18 flat-bottom restraints had a force constant of 5 kcal/mol/Å^2^, with no force applied until the C_α_ distance deviated more than ±2 Å from the distances observed in the experimental structure. These distance restraints were applied to maintain an open gate at the narrowest point. Two identical simulations were also run starting with the low-K structure for 0.5 μs each. In this case, an equivalent constraint was applied to maintain the identified H bond between the S620 hydroxyl H atom and the flipped carbonyl O atom of residue V625.

#### Umbrella Sampling simulations of the V625 carbonyl

Umbrella sampling simulations^70^ of the rotation of the V625 carbonyl was performed by constraining the V625 backbone 4¢¢ angle (N_625_-C_625_-Ca_625_-N_626_). The backbone 4¢¢ angle controls the V625-Gly626 linkage and thus the orientation of the V625 carbonyl oxygen (with 4¢¢∼-50° leading to the carbonyl oxygen pointing in and 4¢¢∼+70° leading to the carbonyl oxygen pointing out). This backbone dynamics has previously been shown in both KcsA and MthK^15,30^. Initial windows were created using steered MD with a harmonic force constant of 0.03 kcal/mol/°^2^ moving at a rate of 0.2 ns/°. The complete 360 ° 4¢¢ rotation was divided into 72 windows separated by 5°. The backbone dihedral was constrained with a force constant of 0.03 kcal/mol/°^2^ in the center of each window. During the production run, multi-ions configurations in sites S0/S2/S4, S2/S4, S2/S3 and S1/S3 were constrained with 5 kcal/mol in their sites with flat-bottom potentials (k=10 kcal/mol/Å^2^ and width 3.5 Å). Simulations were performed for the WT channel with ions in S0/S2/S4, S1/S3. A convergence criterion of a free energy change of less than 1 kcal/mol was used and this was achieved after 9 ns and 11 ns for WT with ions in S0/S2/S4 or S1/S3. All data prior to equilibration was discarded for final calculations. WHAM^71^ with periodic boundary conditions was used to resemble and unbiased the free energy profile. Error bars were calculated as standard error of mean by dividing the data into 1 ns long blocks.

#### Replica Exchange with Solute Tempering

To study the behavior of the filter at 300K temperature, replica exchange with solute tempering (REST2) simulations^32^ starting with the conductive structure were performed. 3 K^+^ ions were trapped in the vicinity of the selectivity filter/cavity region using a tall cylinder of 30 Å height and 15 Å width with flat-bottom half-harmonic constraints of 10 kcal/mol/Å^2^. 16 replicas with effective temperatures for ion interactions from 300 K to 900 K were used, where only the interactions involving the 3 trapped ions were scaled. Two independent systems, with ions starting in S0/S2/S4, or with S1/S3/cavity, were simulated for 500 ns each. The first 14 ns were removed from each simulation before calculating the final free energy profile as Δ*G*(*z*) = −*k_B_T* ln ρ(*z*) + *C*. Where *ρ* is the probability distribution as a function of reaction coordinate *z*, the position of an ion along the *z* coordinate with respect to the center of mass of the backbone atoms of the selectivity filter and the constant *C*, was chosen to set the free energy to zero in the extracellular solution. This effective free energy profile based on ion density due to the 3 ions trapped inside the filter, reveals the apparent locations and depths of the binding sites, as well as the apparent barriers for ion movement between sites, but is distinct from the barriers that would be seen in a multi-ion permeation mechanism. It is thus used as a guide to understand the propensities for ion binding and movement.

A separate analysis was performed by breaking down the trajectory according to the number of V625 carbonyl oxygens that were pointing towards the channel axis (within ±70 °). Five different “clusters”, with 0, 1, 2, 3 and 4 carbonyl oxygens pointing in, were used, leading to different free energy profiles (Supplementary Fig **S10**), revealing the impact of V625 isomerizations on the free energy of ions in the selectivity filter.

## Acknowledgments

We thank JingTing Zhao and members of the Hill and Vandenberg labs, Michael Clark and members of the Perozo Lab, and Celine Boiteux and members of the Allen Lab for assistance and advice during the project. We thank Terry Campbell, Bob Graham, Richard Harvey, Dan Roden, Benoit Roux, and Mike Sanguinetti for comments on the manuscript. We wish to acknowledge the Victor Chang Cardiac Research Institute Innovation Centre, funded by the NSW Government, and the Electron Microscope Unit at UNSW Sydney, funded in part by the NSW Government. TWA, KMC, BWN, and EF acknowledge support from the Medical Advances Without Animals Trust.

## Funding

Australian Research Council Grants DP150101929 (JIV), DP170101732 (TWA), DP200102540 (JIV, EP), DP210102405 (TWA), DP220103550 (TWA).

National Health and Medical Research Council grants APP1116948 (JIV), App1141974 (JIV, TWA, AGS & EP)

National Computational Initiative dd7 (TWA)

## Author contributions

JIV, TWA & EP designed and conceived the study. CL, MJH & JIV designed and performed biochemistry experiments. CL, MJH, AGS, JB, EP & JIV designed and performed cryo-EM experiments; CL, MJH, AGS, EP & JIV analyzed cryo-EM datasets. TWA, EF & JIV designed the simulation experiments; EF, KMC, BWN & TWA performed and analyzed the simulations. MJH, C-AN & JIV designed and performed electrophysiology experiments. The manuscript was written by CL, EF, MJH, EP, TWA & JIV with input from all authors.

## Competing interests

The authors declare no competing interests.

## Inclusion and Diversity

We support inclusive, diverse, and equitable conduct of research.

## Data and materials availability

Maps and atomic co-ordinates for the structures are available at the Electron Microscopy Data Bank (https://www.ebi.ac.uk/pdbe/emdb/) and the Protein Data Bank (http://www.rcsb.org). WT High-K C4: PDB: 8U67, EMD-41955; WT low-K C4: PDB:8U68, EMD-41958

## Supplementary Materials

Supplementary Table S1

Supplementary Figures S1 to S12

Supplementary Movie S1

## Notes

### Competing Interest Statement

The authors have declared no competing interest.

