## Supplementary Tables and Figures for "Potassium dependent structural changes in the selectivity filter of HERG potassium channels"

#### Supplementary Material

Table S1: Cryo-EM data collection, refinement, and validation statistics

Figure S1: Cryo-EM workflow

Figure S2: TMD densities for high-K and low-K

Figure S3: Ramachandran plots for high-K and low-K pdb

Figure S4: MDFF fits to cryo-EM maps for high and low-K structures.

Figure S5: Hole profiles of HERG high- and low-K compared to KcsA.

Figure S6: Ion conduction events in HERG

Figure S7: Constrained ion MD simulations to explore the effect of $K^+$ ions on selectivity filter conformation.

Figure S8: REST2 simulations of HERG

Figure S9: Cluster analysis of constrained ion MD simulations

Figure S10: Interactions behind the selectivity filter for MD simulations with different $K^+$ ion conformations.

Figure S11: Changes in water distribution behind the selectivity filter

Figure S12: 2-dimensional free energy maps for V625 carbonyl oxygen flipping

Movie S1: Transition between conducting and non-conducting states

#### Cryo-EM data collection, refinement and validation statistics*

| | 300 mM $K^+$ C4 symmetry (EMDB-41955) (PDB 8U67) | 300 mM $K^+$ C1 symmetry (EMDB-43006) (PDB 8V7M) | 3 mM $K^+$ C4 symmetry (EMDB-41958) (PDB 8U68) | 3 mM $K^+$ C1 symmetry (EMDB-43007) (PDB 8V7N) |
|---|---|---|---|---|
| **Data collection and processing** | | | | |
| Magnification | 130,000 x | | 130,000 x | |
| Voltage (kV) | 300 | | 300 | |
| Electron exposure (e–/Å$^2$) | 50 | | 50 | |
| Defocus range (μm) | -0.6 to -2.5 | | -0.6 to -2.5 | |
| Pixel size (Å) | 1.1 | | 1.1 | |
| Symmetry imposed | C4 | C1 | C4 | C1 |
| Initial particle images (no.) | | | | |
| Final particle images (no.) | 232,371 | | 358,026 | |
| Map resolution (Å) | 3.3 | 3.5 | 3.0 | 3.4 |
| FSC threshold | 0.143 | 0.143 | 0.143 | 0.143 |
| Map resolution range (Å) | 3.2 – 5.4 | 3.4 – 5.9 | 2.9 – 4.9 | 3.3 – 5.5 |
| **Refinement** | | | | |
| Initial model used | 5VA1 | This study | 5VA1 | This study |
| Model resolution (Å) | 3.4 | 3.8 | 3.1 | 3.4 |
| FSC threshold | 0.5 | 0.5 | 0.5 | 0.5 |
| Model resolution range (Å) | | | | |
| Map sharpening *B* factor (Å$^2$) | -108.0 | -120.1 | -86.6 | -105.4 |
| Model composition | | | | |
| Non-hydrogen atoms | 9508 | | 9508 | |
| Protein residues | 1172 | | 1172 | |
| Ligands | 0 | | 0 | |
| *B* factors (Å$^2$) | | | | |
| Protein | 46.01 | 47.16 | 70.03 | 82.54 |
| Ligand | n.a. | n.a. | n.a. | n.a. |
| R.m.s. deviations | | | | |
| Bond lengths (Å) | 0.013 | 0.012 | 0.013 | 0.012 |
| Bond angles (°) | 1.747 | 1.867 | 1.730 | 1.862 |
| Validation | | | | |
| MolProbity score | 0.5 | 0.5 | 0.5 | 0.53 |
| Clashscore | 0.00 | 0.00 | 0.00 | 0.00 |
| Poor rotamers (%) | 0.00 | 0.00 | 0.00 | 0.00 |
| Ramachandran plot | | | | |
| Favored (%) | 98.25 | 98.07 | 98.6 | 97.89 |
| Allowed (%) | 1.75 | 1.93 | 1.4 | 2.11 |
| Disallowed (%) | 0.0 | 0.0 | 0.0 | 0.0 |

### Supplementary Fig S1

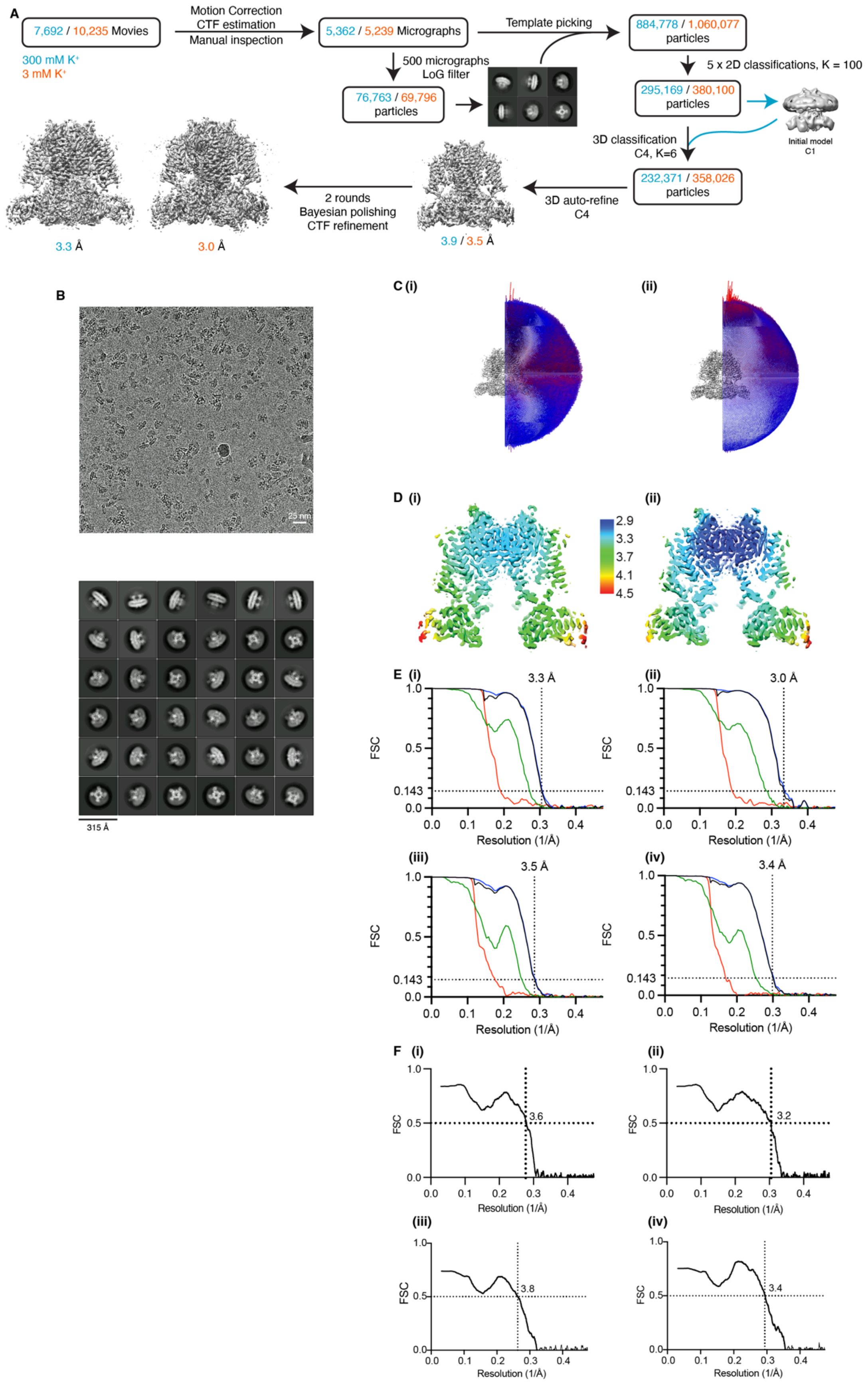

**Supplementary Fig 1**: **Summary for cryo-EM analysis of hERG Ts.**

**A.** Flow chart of EM data processing for high-K (cyan) and low-K (orange) datasets. Details can be found in Methods section. **B.** Representative electron micrograph and 2D class averages of hERG Ts for low-K dataset. **C.** Euler angle distribution of all particles used for final 3D reconstruction for (i) high-K and (ii) low-K dataset. **D.** Local resolution projected on a centre cross-section of the final 3D reconstruction. **E.** Gold-standard Fourier shell correlation (FSC) curve for the 3D reconstruction for (i) high-K with C4 symmetry, (ii) low-K with C4 symmetry, (iii) high-K with C1 symmetry, (iv) low-K with C1 symmetry. **F.** Gold-standard Fourier shell correlation (FSC) curve between the map and model for (i) high-K with C4 symmetry, (ii) low-K with C4 symmetry, (iii) high-K with C1 symmetry, (iv) low-K with C1 symmetry.

#### Supplementary Fig S2

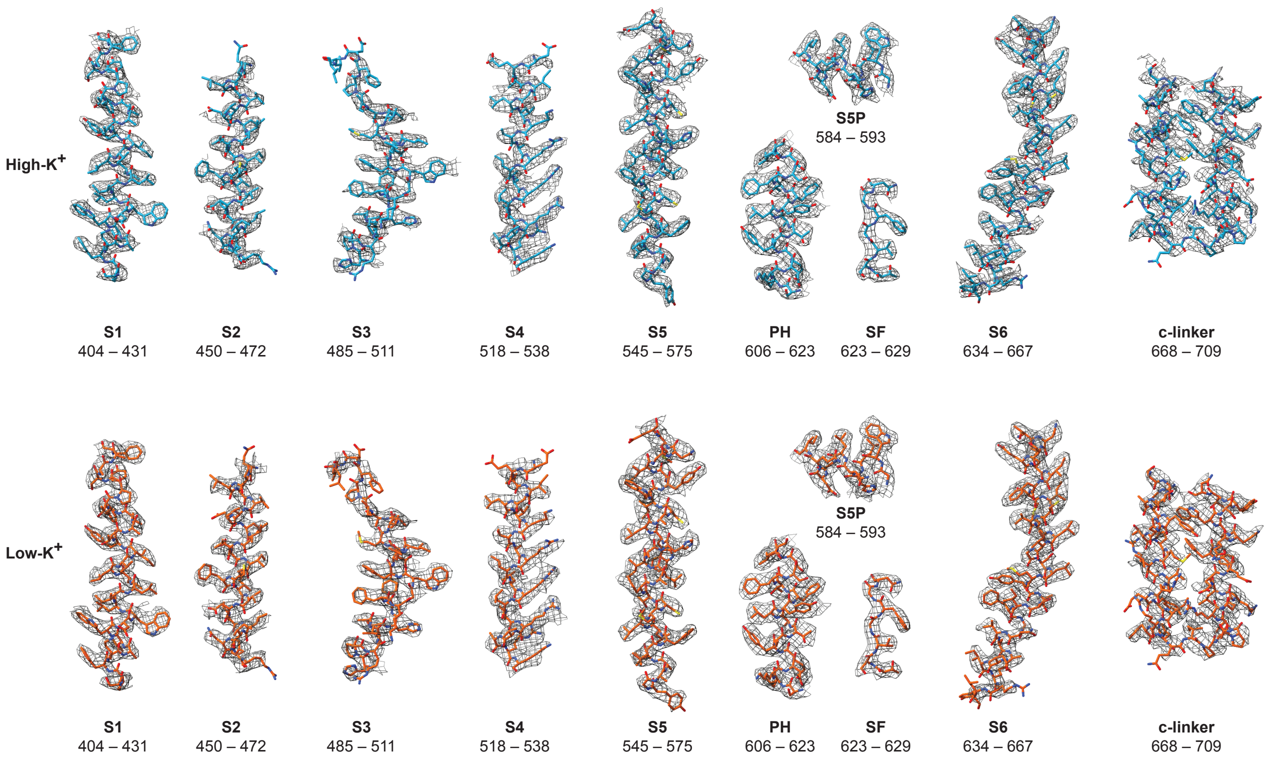


**Supplementary Fig 2: Cryo-EM density maps of selected regions of WT HERG in high-K (300 mM KCl) and low-K (3 mM KCl)**

Atomic models are shown as blue (high-K) and orange (low-K) sticks. Density maps are shown as grey wire mesh. Maps were sharpened using b-factor listed in Table S1.

#### Supplementary Fig S3

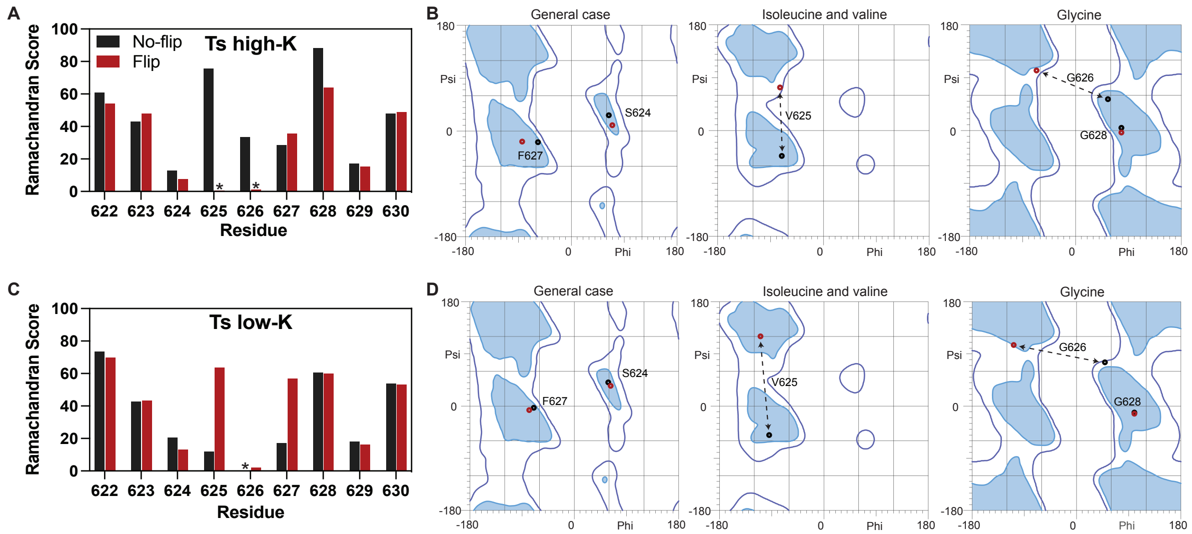


**Supplementary Fig 3: Comparison of Ramachandran statistics for flipped and non-flipped V625 carbonyl oxygen in high-K and low-K model**

**A.** Summary of Ramachandran probability for flipped and non-flipped V625 carbonyl oxygen in high-K models. **B.** Ramachandran plots for residues 624–628 of flipped and non-flipped high-K model. **C.** Summary of Ramachandran probability for flipped and non-flipped V625 carbonyl oxygen in low-K models. **D.** Ramachandran plots for residues 624–628 of flipped and non-flipped low-K models.

**Supplementary Fig S4**

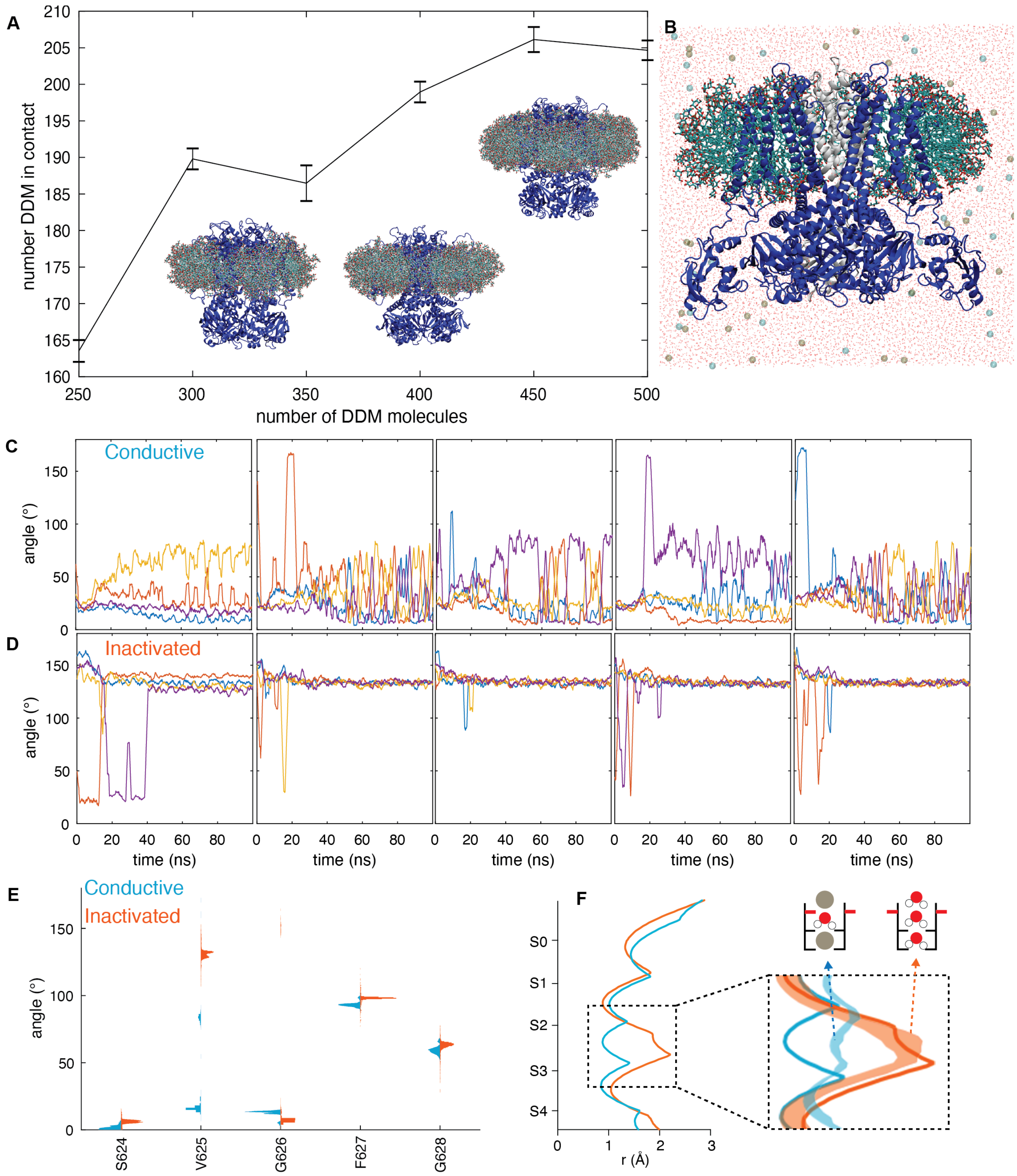


**Supplementary Fig 4 : MDFF fits to cryo-EM maps for high and low-K structures.**

**A**. The optimal size of the micelle was determined by simulating HERG (PDB:5VA1) in 6 different sizes between 250 and 500 molecules of a simple pure detergent micelle (DDM). The number detergent molecules in contact with the protein is plateauing at 400-450 molecules showing that this is the

(Legend continued over page)

(shown in blue, with the TMD of the subunit at the front hidden and at the back in grey, for clarity) embedded in a micelle of 404 molecules consisting of detergent (DDM (300)), lipids (POPE (20), POPC (20), POPA (4)) and cholesterol derivative (cholesteryl hemisuccinate (60)) (shown as cyan (C), blue, (N), red (O) and olive (P) sticks) and surrounded by TIP3P water molecules[56] (shown as red sticks) and KCl (shown as olive and cyan spheres, respectively). **C-D.** Timeseries of the V625 carbonyl oxygen angle for the five independent MDFF simulations for high K channel with ions in S2/S4 (**C**) and low-K channel with an empty selectivity filter (**D**). For both systems, we see flipping and unflipping in the early part of the simulations. The high K channel converges to a mix of flipped and unflipped V625 carbonyl oxygens, whereas in the low-K channel all V625 are flipped at the end of the simulations. **E**. Distribution plots of the angle of the backbone carbonyl oxygens in the selectivity filter for the high (blue) and low (orange) $K^+$ structures based on 250 energy minimised structures extracted from MDFF simulations. **F**. Plot of the selectivity filter radius for the high K (blue) and low-K (orange) channel determined with HOLE[17]. Solid lines show the radius determined from the cryo-EM fit and shaded area the radius determined for the 250 energy minimised structures extracted from MDFF simulations. The decreased radius in the MDFF simulations compared to the cryo-EM fit at the V625 carbonyl oxygens (between S2 and S3) in the high K channel arise comes from the V625 carbonyl oxygens being in a mix of flipped and non-flipped states.

**Supplementary Fig S5**

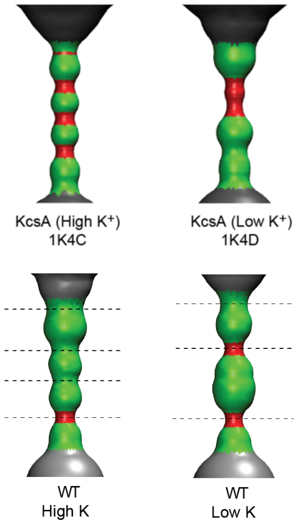


**Supplementary Fig 5: HOLE profiles for High-K and low-K HERG compared to High-K and low-K KcsA**

Shape of ion conduction pathway calculated using HOLE. All models are aligned to the pore helix. The KcsA low-K profile is quite distinct to the HERG low-K structure, suggesting there may be different inactivation mechanism for the two channels.

**Supplementary Fig 6**

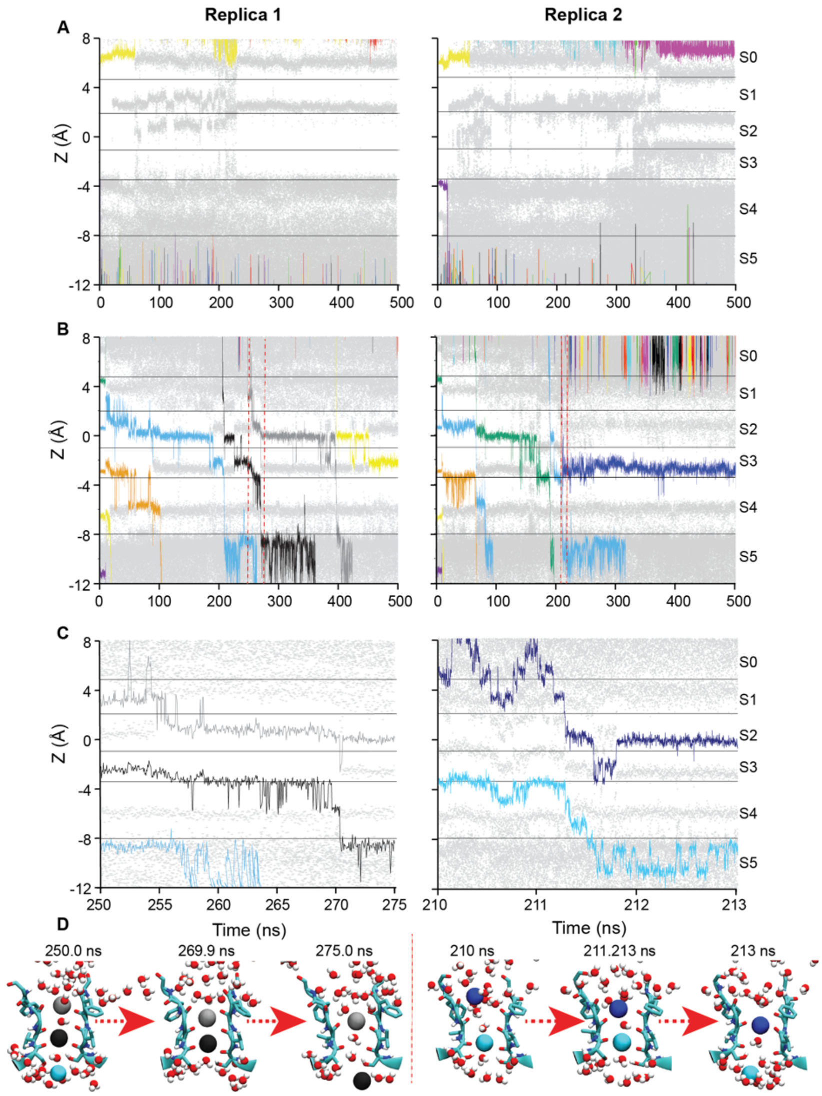


**Supplementary Fig 6: Ion conduction in HERG structures**

Replica simulations for **A.** Low-K filter and **B.** High-K filter in the presence of a 500 mV gradient (inwards negative). Individual potassium ions are shown as colored lines, water molecules are shown as grey dots. In the low-K filter a loose constraint (see methods) was imposed to maintain the V625 carbonyl oxygens pointing outwards and in the high-K filter a loose constraint (see methods) was imposed to maintain the V625 carbonyl oxygens pointing inwards and. **A.** There are no conduction events observed when V625 is point outwards. **B.** With the V625 pointing inwards there are multiple ion-water-ion mediated (soft knock-on) conduction events. In the right-hand panel, there are no conduction events after 225 ns, which correlates with an outward rotation (~90°) of the F627 carbonyl oxygens and the presence of more water molecules in the upper filter. **C.** Expanded trace showing (i) the only example we observed of a hard knock on (ion-ion) event and (ii) an example of a soft knock-on (ion-water-ion) event. **D.** Structure snapshots taken at the times indicated to illustrate the ion conduction events highlighted in **C**.

**Supplementary Fig 7**

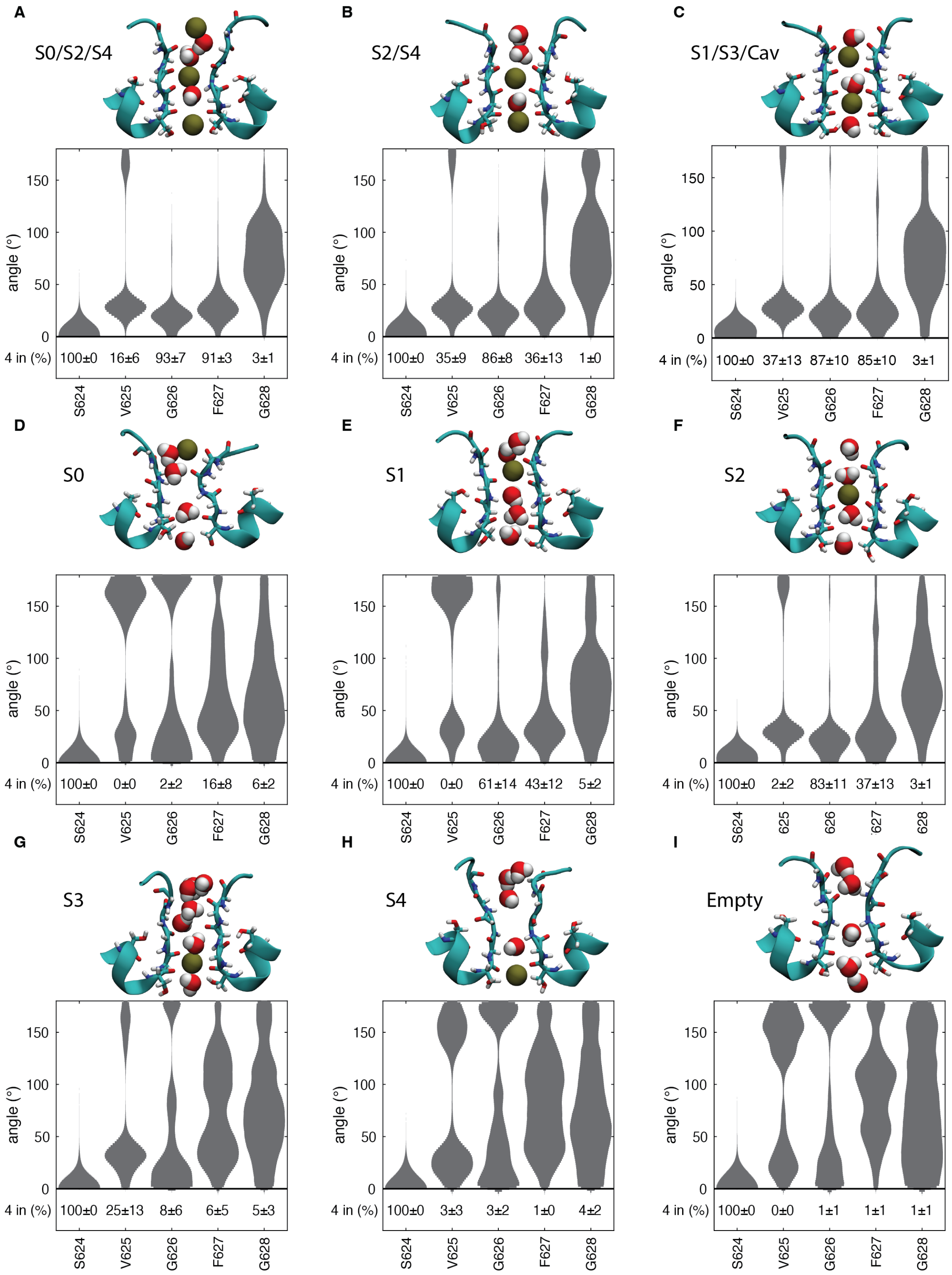


See overleaf for legend.

**Supplementary Fig 7: MD simulations to explore the effect of $K^+$ ions on selectivity filter conformation.**

Distribution plots of the angle of the backbone carbonyl oxygens in the selectivity filter (defined as the absolute value of the angle between the vector made up by Cα and the carbonyl oxygen atom and Cα and the center of mass of the selectivity filter) (top) and the frequency of four carbonyl oxygens pointing in (with an angle of < 70 degrees) (bottom) for ions held in **A.** S0/S2/S4, **B.** S1/S3/Cav, **C.** S2/S4, **D.** S0, **E.** S1, **F.** S2, **G.** S3, **H.** S4, and **I.** an empty selectivity filter. Insets in each panel show typical ion configurations. When $K^+$ ions were maintained in S0/S2/S4, S2/S4 or S1/S3/Cav (**A-C**), there were periods of time when V625 carbonyl oxygens were seen to flip to no longer be directed into the pore, with all four carbonyl oxygens pointing towards the channel axis only 16%, 35% and 37% of the time. However, when the filter was empty (**I**), there were never four V625 carbonyl oxygens pointing towards the channel axis. When an ion was placed in S0, S1 or S4 (**D,E,H**), the selectivity filter resembles that of the empty filter, with two or more carbonyl oxygens flipped 95%, 99% and 56% of the time, respectively, shown by a larger density of big angles in the probability like that of an empty selectivity filter (**I**). When an ion was placed in S2 or S3 (f and g), the selectivity filter resembles that of a filter with a multiple-ion configuration (carbonyl oxygens mostly directed inward with three or more carbonyl oxygens pointing in 94% and 80% of the time) (**A-C**). Even with an ion held in S2, we still observed that V625 can flip, with one or two flipping 91% and 6% of the time, respectively.

**Supplementary Fig 8**

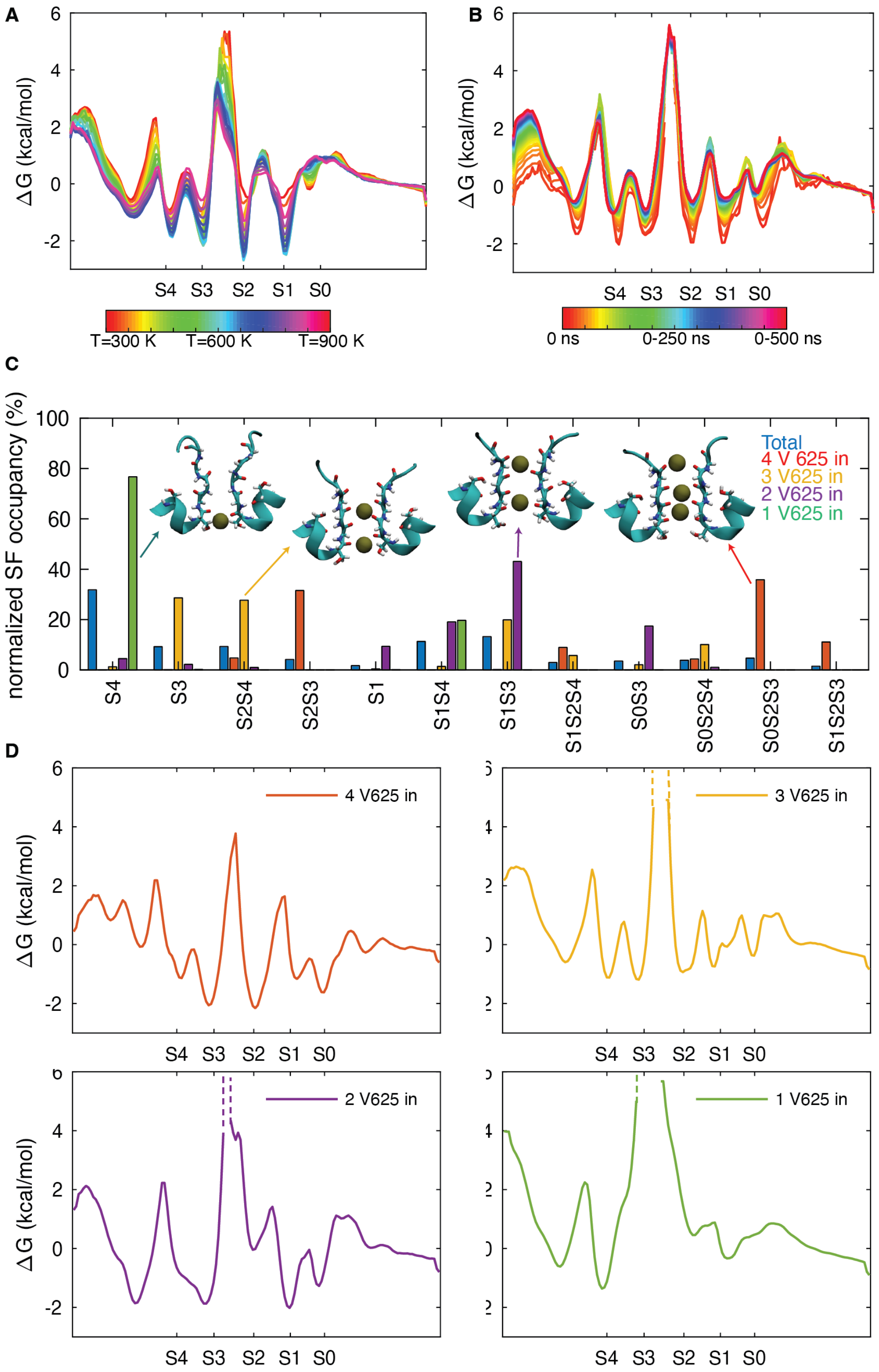


Legend: see overleaf

**Supplementary Fig 8: REST2 simulations**

Free energy profile for all 16 replicas with temperatures varying from 300 K to 900 K for REST2 simulations[22] with ions starting in **A.** S0/S2/S4 and **B.** S1/S3/Cav. The higher temperature replicas show shallower minima and lower barriers with a well captured barrier between S2 and S3. The free energy profile shows deepest binding in S0, S2 and S3 with S1 and S4 elevated by 1 kcal/mol. There are small barriers for movement between S0-S1 and S3-S4 (~1 kcal/mol) with medium barriers for movement between S1-S2 and to exit the filter (~2-3 kcal/mol), and a large barrier (~6 kcal/mol) for crossing between S2-S3. **C-D.** Throughout the REST2 simulations, we observed extensive fluctuations in the selectivity filter, with frequent flipping and unflipping of V625 carbonyl oxygens. When we separated the data into bins according to the number of flipped V625 carbonyl oxygens it becomes hard to sample the high barrier between S2 and S3, when only a small fraction of the trajectory is included, and so it is only well defined when all four V625 carbonyl oxygens point inwards. When all four V625 carbonyl oxygens were directed inward (13 % of the time; shown in red), there was on average 2.7 ions in the selectivity filter, most commonly with ions in S2/S3, S0/S2/S3 or S1/S2/S3. More commonly, when three V625 carbonyl oxygens were pointing in towards the channel axis (31% of the time; shown in yellow), there was an average of 1.9 ions in the selectivity filter maintaining good binding in all sites. In the free energy profile, a displacement of the S3 ion downwards can be seen and S1 and S2 are not well defined, instead the free energy profile shows multiple minima in this area (e; yellow). These arise when the Tyr627 and Gly628 carbonyl oxygens flip and the selectivity filter widens, which creates off axis binding where the ion interacts with four carbonyl oxygens from two neighbouring residues and subunits. When two carbonyl oxygens were pointing inwards (16% of the time, almost exclusively in neighbouring subunits; shown in purple) there was, on average, 2.0 ions in the selectivity filter, most commonly occupying S3 as well as S1 or S2 (purple). S3 and S4 have now merged into one site which is positioned around the Ser624 carbonyl oxygen plane. Noticeably, there is reduced binding in the S2 site that may be expected to inhibit conduction. 39% of the time a single carbonyl oxygen was pointing in towards the channel axis (shown in green), with an average ion occupancy of 1.5. Now the ions almost exclusively bind to S4 and on the G626 carbonyl oxygen plane between S0 and S1, vacating the entire centre of the selectivity filter and prohibiting ion translocation. It was very unlikely (0.0052% of the time) that all V625 carbonyl oxygens were flipped outward (not shown).

#### Supplementary Fig 9

A

| | Cluster occupancy (%) | | | | | | | |
|---|---|---|---|---|---|---|---|---|
| Cluster | (1) Conductive-like | (2) Inactivated-like | (3) Flipped G626 | (4) Flipped F627 | (5) Flipped V625 and G626 | (6) Flipped F627 | (7) Flipped G626, F627 and G628 | (8) Flipped G628 |
| total | 41 | 30 | 4.8 | 4.8 | 4.7 | 4.7 | 4.2 | 3.6 |
| S0/S2/S4 | 73 | 22 | 0 | 0.61 | 0.025 | 0.062 | 0 | 3.8 |
| S2/S4 | 64 | 17 | 0 | 1.1 | 0.055 | 8.9 | 0 | 8.2 |
| S1/S3/Cav | 71 | 17 | 2.4 | 2.2 | 0.33 | 1.4 | 0.14 | 5.7 |
| S0 | 11 | 41 | 9.8 | 9.7 | 16 | 2.9 | 4 | 1.7 |
| S1 | 25 | 53 | 4.4 | 8.3 | 3.8 | 3.7 | 0.73 | 1.2 |
| S2 | 57 | 25 | 2.5 | 4.2 | 0.34 | 4.1 | 0.20 | 4.2 |
| S3 | 45 | 21 | 6.1 | 2.9 | 0.054 | 10 | 6.3 | 4.0 |
| S4 | 19 | 33 | 13 | 7.1 | 5.5 | 8.2 | 12 | 0.14 |
| Empty | 7.5 | 39 | 5.5 | 7.2 | 16 | 2.5 | 14 | 3 |

B

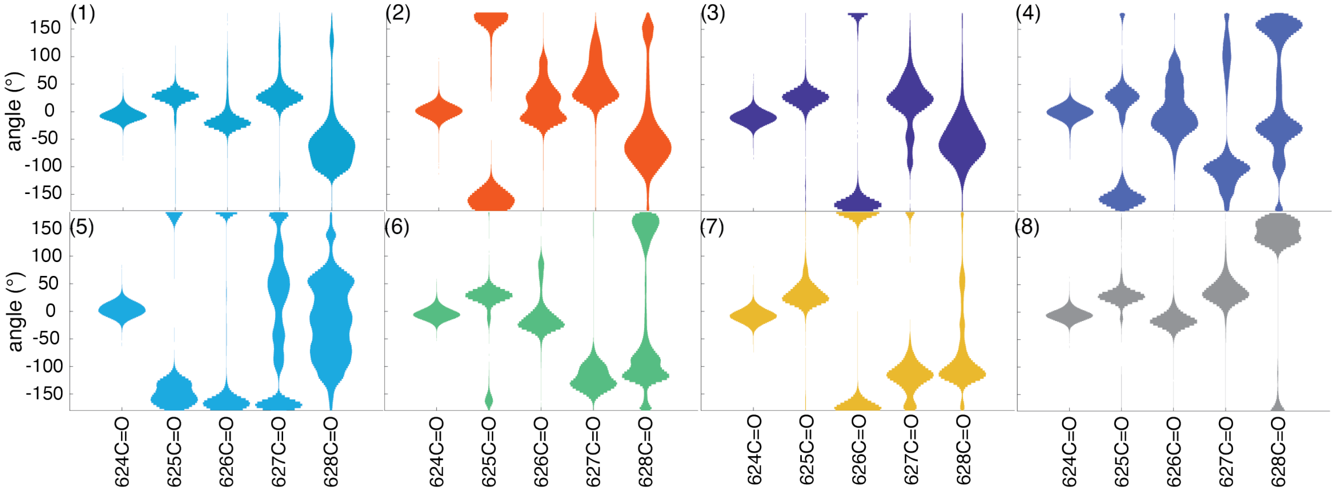


**Supplementary Fig 9: MD simulations with different $K^+$ ion conformations clustered according to backbone dihedral angles of each subunit of the selectivity filter.**

**A.** Table showing the occupancy (rows) for the 8 major clusters (labelled 1-8 in **A** and **B**) obtained from analysis of Selectivity Filter backbone atoms across all MD simulations (columns). The occupancy in each cluster across all simulations is shown on row 3 and for each ion-configuration simulation on rows 4-12. Cluster (1) shows a conductive-like state with the V625 carbonyl oxygen pointing towards the channel axis (41% total occupancy) and cluster (2) shows an inactivated-like state with the V625 carbonyl oxygen pointing out with a narrowing at G626 (30% total occupancy). Analysis of these two clusters are shown in Figure 3 in the main manuscript. The additional 6 clusters, which had various backbone residues flipped, had occupancies between 3% and 5. When ions were placed in S0/S2/S4, S2/S4, S1/S3 or S2, the subunit maintained a conductive conformation most of the time (a; second column). This suggests that there must be multiple ions in the selectivity filter (including S2 or S3), or S2 must be occupied, for the selectivity filter to be conductive. **B.** Distribution plots of the angle of the backbone carbonyl oxygens in the selectivity filter for the 8 most observed clusters.

**Supplementary Fig 10**

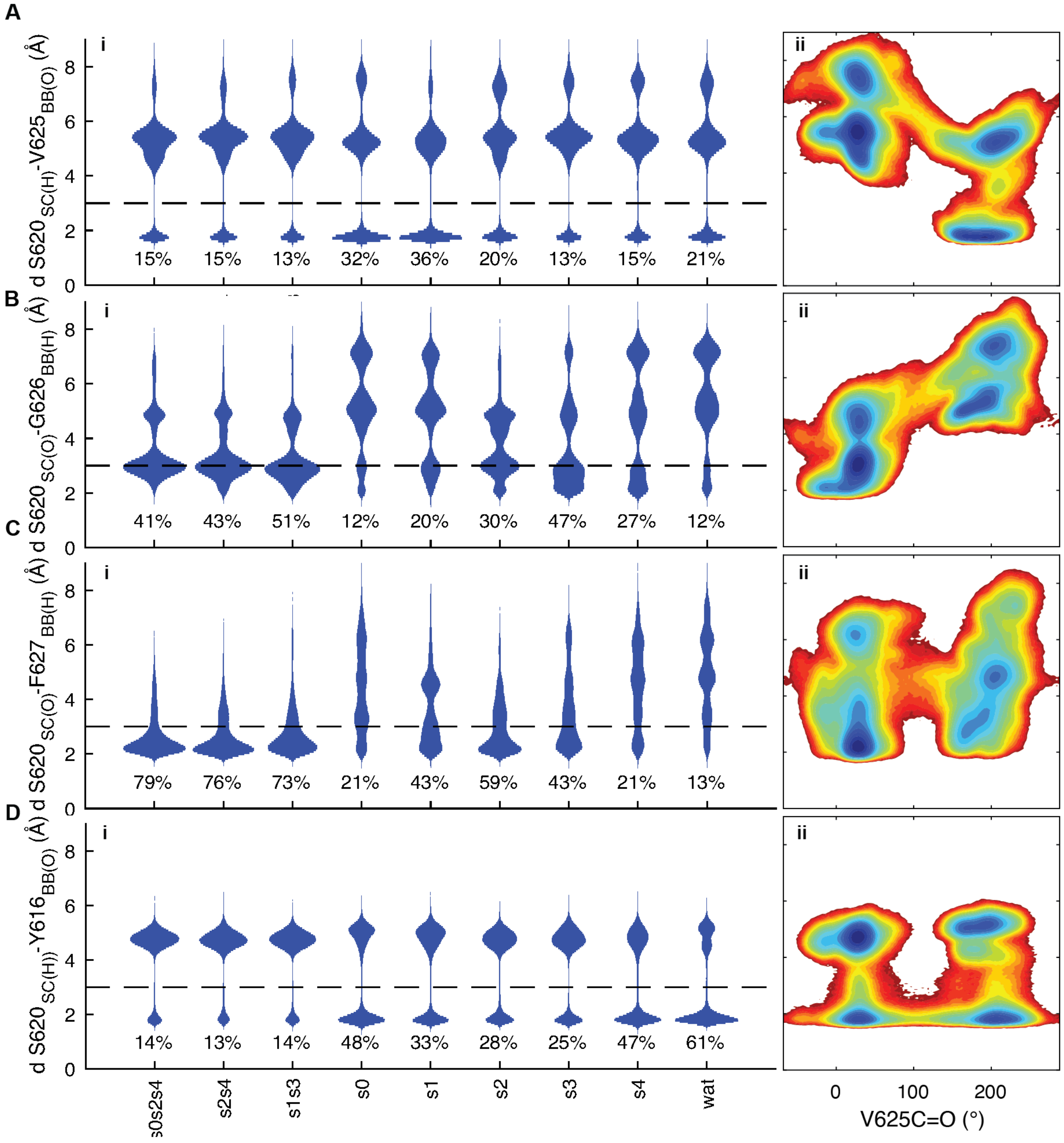


**Supplementary Fig 10: Interactions behind the selectivity filter for MD simulations with different $K^+$ ion conformations.**

**A-D.** Distribution plots for the distance between the S620 sidechain hydroxyl and V625 C=O (**A**), G626 N-H (**B**), F627 NH (**C**) and Y616 C=O (**D**), as a function of ion occupancy of the filter (i) and 2-dimensional free energy plots as a function of rotation of the V625 carbonyl oxygen (ii). The 2D FEP obtained from the constrained ion MD, shown here, are very similar to what we observed with the REST2 simulations (see Figure 4D-G in main text).

**Supplementary Fig 11**

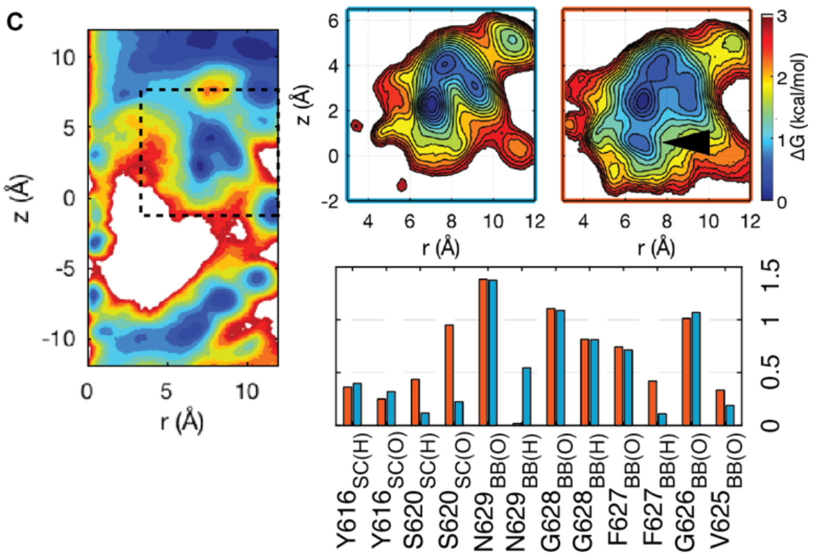


**Supplementary Fig 11: Changes in water distribution behind the selectivity filter in Cluster 1 (conducting) and Cluster 2 (non-conducting) states of HERG**

**A.** Free energy distribution of number of water molecules behind the filter in all simulations. **B.** Free energy distribution of water molecules behind the filter in (i) conducting state (cluster 1, Supplementary Fig 9) and (ii) non-conducting state (cluster 2, Supplementary Fig 9). Arrowhead highlights an extra minimum in the non-conducting state. **C.** Mean number of interactions with water molecules behind the selectivity filter in conducting (blue) and non-conducting (orange) filters.

The additional minimum in the non-conducting state corresponds to a water molecule that predominantly interacts with the S620 side chain. This site is similar in location to the deepest water molecule in the inactivated KcsA structure although in KcsA that water binds to and stabilizes the pinched glycine amide[17]. There is also an increase in water bound to the Phe627 backbone amide when the V625 carbonyl oxygen is flipped, that compensates for F627 no longer interacting with S620 (**C**). It is notable that the total number of water molecules is not different between the flipped and non-flipped V625 carbonyl oxygen states of HERG (**C**), which is in marked contrast to KcsA, where the number of water molecules increases from one in the conductive state to three in the non-conductive state and this diffusion-limited transition plays a key role in establishing the slower inactivation kinetics of KcsA[17].

#### Supplementary Fig 12

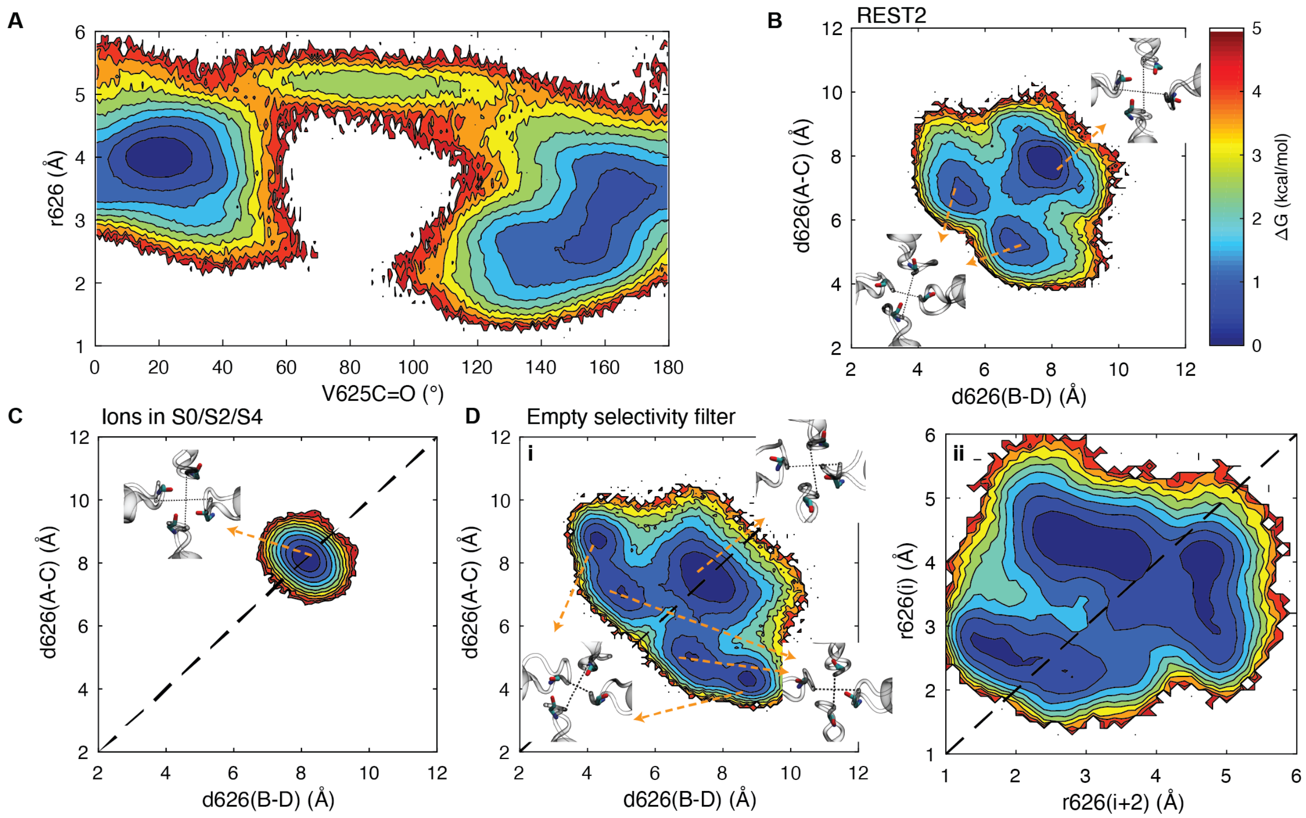


**Supplementary Fig 12: 2-dimensional free energy maps for V625 carbonyl oxygen flipping**

**A.** 2-dimensional free energy maps highlighting the relationship between the flipping of the V625 carbonyl oxygen (>70° away from the central axis) and the pinching of the G626 (represented as a mean radial position of G626 Cα atoms, r626) . **B-D.** 2-dimensional free energy maps showing the pinching of G626 Cα based on the distance between opposing subunits in: REST2 simulations **B**; MD simulations with ions constrained in S0/S2/S4; **C,** and for an empty selectivity filter **(Di)**. **Dii.** 2-dimensional free energy map showing the relationship between the radial positions of G626 Cα in two opposing subunits during the pinching of G626. Although **Di** suggests an asymmetric collapse with 2-fold symmetry, it is apparent from **dii** that this arises predominantly due to movements only in one of the two subunits, which contrasts with the C2 symmetrical inactivated state suggested by Li and colleagues[23] .

**Supplementary Movie 1**

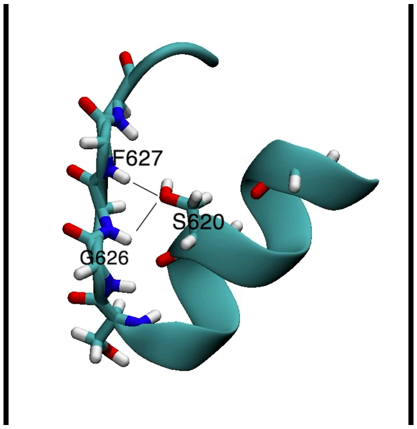


**Supplementary Movie 1. Interactions between S620 and the selectivity filter during the transition between conducting and non-conducting structures**

Movie showing how the S620 sidechain can interact with the backbone of G626 and F627 (H) or the backbone of Y616 (O) in the conductive state (Figure 4 D-G (i), main text). When the sidechain of r620 interacts with the backbone of Y616 (O) ((Figure 4 D-G (ii),(iii) main text), the V625 carbonyl oxygen can flip outward. The sidechain of S620 can then interact with the backbone of V625 (O) (Figure 4 D-G (iv), main text) in the inactivated state.
